# Multi-Scale Anti-Correlated Neural States Dominate Naturalistic Whole-Brain Activity

**DOI:** 10.1101/2025.08.27.672600

**Authors:** Dora Gözükara, Djamari Oetringer, Nasir Ahmad, Linda Geerligs

**Author notes:** **For correspondence:** (DG). Denotes shared senior authorship (Nasir Ahmad, Linda Geerligs). Thomas Van Aquinostraat 4, 6525GD Nijmegen, Netherlands.

## Abstract

The human brain’s response to naturalistic stimuli is characterized by complex spatiotemporal dynamics. Within these dynamics there is a transitioning structure between sets of anti-correlated neural states that is frequently observed but has not been systematically investigated across scales. In this paper, we use three different naturalistic fMRI datasets to quantify anti-correlation in global and local neural states during naturalistic viewing or listening and investigate their interdependence and their relationship to changes in the stimuli. We demonstrate that continuous naturalistic brain activity shows an anti-correlational structure that spans both global and local spatial scales, with regions in the dorsal attention network showing strong alignment between local and global state transitions. On the global scale, ongoing dynamics are dominated by two antagonistic states that correspond to Default Mode Network and Task Positive Network configurations, with a third transitional state mediating between them. On the local scale, we observe anti-correlated neural states that are associated with periods of relatively high and low brain activity. Across the brain, these are driven by subsets of voxels that are systematically anti-correlated with their area’s dominant activity pattern. This antagonism is related to stimulus changes, which tend to trigger a switch to the TPN state globally and to high activity states locally. On the local scale we also see a modality-specific pattern, with visual changes mostly driving transitions in visual cortical regions and auditory changes predominantly affecting auditory and language-related areas. The consistency of these findings across datasets with different stimulus types (audiovisual and purely auditory) indicates that anti-correlated neural states represent a domain-general organizational principle of brain function. We propose that anti-correlated dynamics functionally represent a convergent solution to the fundamental challenge of maintaining coherent internal representations while remaining responsive to meaningful changes in the environment.

## Introduction

During naturalistic perception, such as movie watching or story listening, the brain must process continuous sensory input while maintaining contextual information over extended timescales. This presents a computational challenge: neural systems need to update their representations in response to relevant changes while preserving previous information necessary for ongoing comprehension. Understanding how the brain achieves this balance remains an open question in neuroscience.

Literature has long established that continuous brain activity organizes into discrete states that persist for extended periods before transitioning rapidly to new configurations (***Baldassano et al., 2017***; ***Geerligs et al., 2022***; ***Abbas et al., 2019***; ***Yamashita et al., 2021***; ***Abeles et al., 1995***). An intriguing aspect of these dynamics is the presence of anti-correlated patterns—both in large-scale networks and local brain regions—where certain activity configurations alternate with their approximate opposite. These observations suggest that anti-correlated dynamics may play a role in how neural systems balance stability and flexibility during continuous processing. In this study, we systematically examine these anti-correlated patterns across spatial scales during naturalistic viewing to better understand their properties and potential functional significance.

### Neural states in naturalistic processing

The organization of brain activity into discrete states has emerged as a fundamental principle across multiple spatial and temporal scales. During continuous naturalistic stimulation, neural activity patterns remain relatively stable for extended periods—orders of magnitude longer than the transitions between them (***Baldassano et al., 2017***; ***Geerligs et al., 2022***). This phenomenon appears across species, imaging modalities, and cognitive contexts (***Abbas et al., 2019***; ***Yamashita et al., 2021***; ***Abeles et al., 1995***). Neural states also show behavioural relevance across various domains. Global state dynamics have been linked to task performance, attention, narrative comprehension, and exploration-exploitation trade-offs, and even consciousness (***Yamashita et al., 2021***; ***Song et al., 2023***; ***Demertzi et al., 2022***). At finer spatial scales, neural states in different cortical areas are shown to form a partially nested temporal hierarchy, with state transitions shaped by the temporal characteristics of stimulus features relevant to each area’s functional specialization (***Geerligs et al., 2022***; ***Oetringer et al., 2025***). For instance, transitions in the parahippocampal place area align with location changes in narratives, suggesting that local neural states maintain representations of environmentally relevant features. However, accumulating evidence suggests that thinking of neural states as stimulus representations that are extended in time may be an oversimplification. Studies have shown that identical stimuli can evoke anti-correlated neural patterns depending on task relevance (***van Loon et al., 2018***), and careful examination of published data reveals sequential anti-correlation in neural states during naturalistic viewing (see Figure 1 in ***Geerligs et al. (2022***) and Figure 3 in ***Baldassano et al. (2017***)). These observations motivated us to carry a systematic investigation of anti-correlated dynamics in naturalistic settings, across spatial scales.

**Figure 1.**
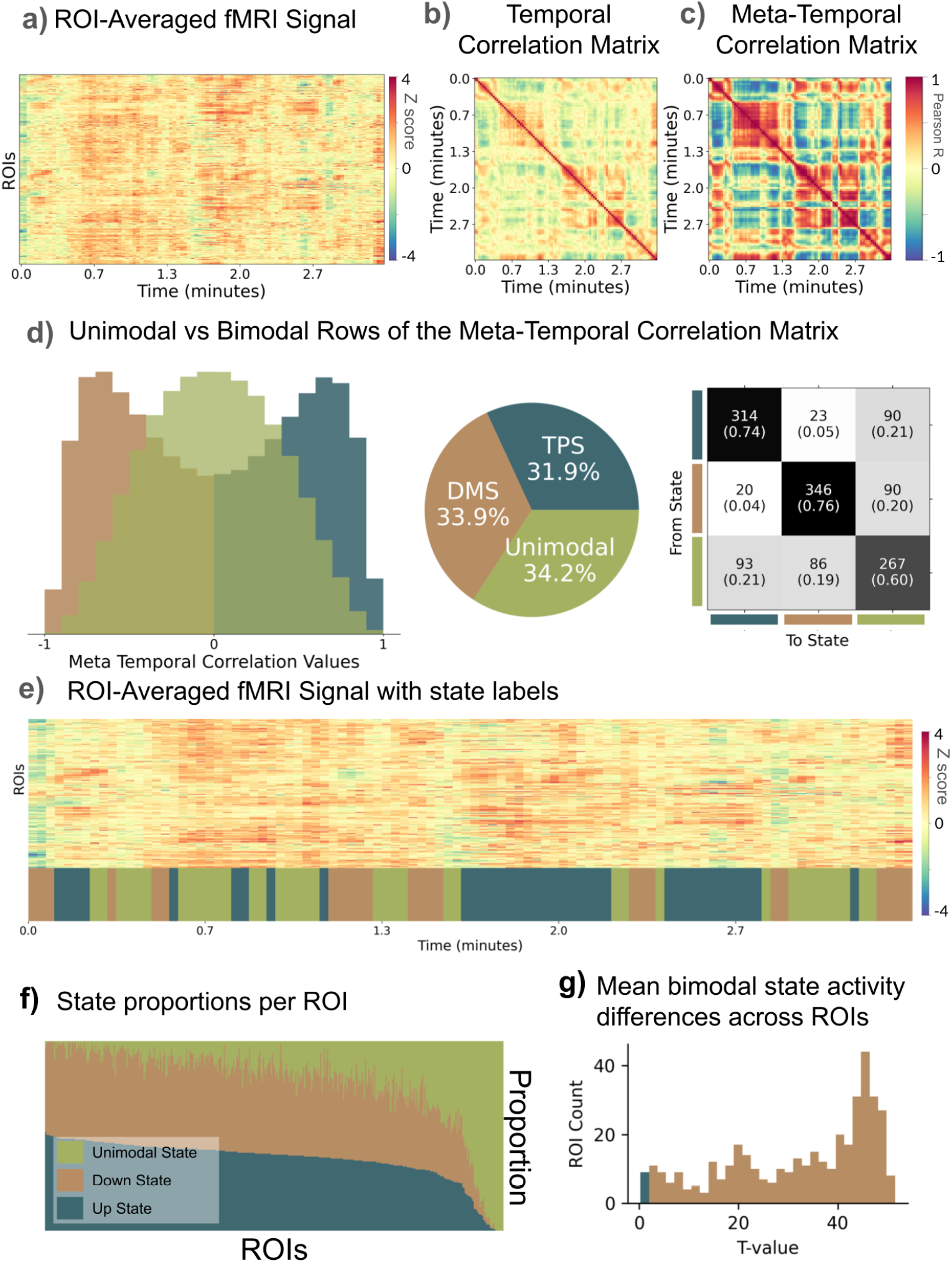
Naturalistic global fMRI activity has an underlying signal which flips between two anti-correlated activity states. (a) Distinct bands appear in the ROI averaged whole-brain time series, implying discrete states. (b) The temporal correlation matrix of the timeseries shows how similar the activity pattern at one timepoint is to all others. It shows prolonged states in which the neural activity patterns are similar over time, which are anti-correlated with the subsequent state. (c) The meta-temporal correlation matrix shows how similar neural patterns at two timepoints are in terms of their relationship to all other timepoints. It shows that the majority of timepoints are either strongly correlated or strongly anti-correlated to each other. (d) Left: The rows of the meta temporal correlation matrix (the relationship of one timepoint to all other timepoints in panel c) are distributed either unimodally or bimodally. Middle: We can thus define three distinct and discrete whole-brain states: Two anti-correlated states and an orthogonal transition state. These three states occupy roughly equal time during naturalistic activity. Right: Transition matrix shows the temporal structure of discrete states. Two anti-correlated states rarely transition to and from one another. A unimodal state mediates the transition from one to the other. (e) Whole-brain timeseries and corresponding brain states. We repeated analyses we conducted on the whole-brain activity for each local ROI (voxel by time) activity timeseries. (f) Local ROIs have fewer unimodal (green) neural states. Each column shows how much of the whole timeseries is spent at each state for an ROI. (g) The mean BOLD activity of one of the bimodal states is consistently higher than its anti-correlational counterpart. Therefore we can call both ends of the local anti-correlated patterns the up and down states. Histogram shows the distribution of t-values resulting from Welch’s t-test comparing the mean activities of the two states across all ROIs. Brown bins show significant t-values and blue bin shows t-values which fall below the significance threshold.

### Global neural states and anti-correlation

At the global scale, brain activity is organized around the well-established antagonistic activation between the default mode network (DMN) and task-positive networks (TPN) (***Fox et al., 2005***; ***Raichle et al., 2001***; ***Greicius et al., 2002***). The DMN, associated with internally directed cognition such as self-reference and memory integration, shows robust anti-correlation with the TPN, which engages during externally focused tasks requiring attention and cognitive control (***Fox et al., 2005***; ***Raichle et al., 2001***; ***Weigard et al., 2024***; ***Cheng et al., 2020***; ***Barber et al., 2016***). This anti-correlational relationship has been extensively documented across resting-state and task-based paradigms, representing a foundational principle of large-scale brain organization (***Fox et al., 2005***; ***Raichle et al., 2001***; ***Greicius et al., 2002***; ***Kelly et al., 2008***; ***Weigard et al., 2024***; ***Barber et al., 2016***).

These anti-correlated dynamics have also been described as antagonistic activity dynamics (***Hammer et al., 2024***; ***Leech et al., 2025***; ***Barber et al., 2016***; ***Cheng et al., 2020***; ***Kucyi et al., 2020***), and quasi-periodic patterns (QPPs)—recurring spatiotemporal templates of activity that alternate between major brain networks (***Majeed et al., 2011***; ***Thompson et al., 2014***; ***Abbas et al., 2019***). These patterns, occurring on timescales of approximately 20 seconds, primarily involve alternating activation and deactivation of the default mode network (DMN) and task-positive network (TPN). Originally discovered in resting state, QPPs have since been observed across various cognitive states and species, suggesting they represent a fundamental organizational principle(***Majeed et al., 2011***; ***Thompson et al., 2014***; ***Abbas et al., 2019***). The behavioural significance of these global anti-correlated states has been demonstrated in several contexts. ***Yamashita et al. (2021***) found that during sustained attention tasks, a DMN-dominant state was associated with higher accuracy and stability, while individuals with ADHD spent more time in a TPN-dominant state. Similarly, ***Song et al. (2023***) linked DMN-dominant states to better narrative comprehension, with transitions between states modulated by global network desynchronization. These findings suggest that the alternation between anti-correlated global states supports adaptive cognitive function. The anti-correlation between DMN and TPN appears to be an active, functionally relevant organization. The DMN, associated with internally-directed processes, and the TPN, engaged during externally-focused attention, show consistent negative correlation across diverse experimental contexts (***Fox et al., 2005***). This relationship aligns with the cortical hierarchy described by ***Margulies et al. (2016***), where the DMN occupies the apex of abstraction while sensory-motor regions anchor the opposite end. The alternation between these anti-correlated states may thus implement a fundamental rhythm between internally and externally directed processing modes.

### Local neural states and anti-correlation

At finer spatial scales, neural dynamics within individual brain regions also organize into discrete states (***Geerligs et al., 2022***; ***Baldassano et al., 2017, 2018***; ***Oetringer et al., 2025***). ***Baldassano et al. (2017***) demonstrated that the brain segments continuous experience into discrete events through shifts in local activity patterns, with different timescales across the cortical hierarchy—shorter states in sensory regions and longer states in high-level areas. This temporal hierarchy provides a multi-scale representation of ongoing experience, from moment-to-moment sensory changes to extended narrative structures. Evidence for anti-correlated patterns within local neural populations comes primarily from task-based studies. Especially in fMRI ***van Loon et al. (2018***) showed that current and prospective goals are represented in opposite patterns within object-selective cortex, suggesting that anti-correlation allows the same neural population to maintain multiple representations without interference. ***Yu et al. (2020a***) extended this finding across brain regions, showing that early visual cortex represents stimulus orientation while intraparietal sulcus represents location through priority-dependent anti-correlated patterns. While these task-based findings demonstrate the capacity for local anti-correlation, whether such patterns occur during naturalistic processing remains unclear. The presence of anti-correlated dynamics at both global and local scales raises the question whether transitions are synchronized across scales and how these transitions relate to naturalistic stimulus processing.

### This study

#### Our study addresses three main research aims

- First, we ask whether anti-correlated patterns are a dominant feature of neural activity during movie watching and story listening. While anti-correlated patterns have been documented in both resting state and task-based studies (***Fox et al., 2005***; ***Raichle et al., 2001***; ***Weigard et al., 2024***; ***Cheng et al., 2020***; ***Barber et al., 2016***; ***Hammer et al., 2024***; ***Leech et al., 2025***; ***Kucyi et al., 2020***), their prevalence and organization during naturalistic processing remains incompletely characterized.
- Second, we investigate how anti-correlated dynamics at global and local scales relate to each other temporally and spatially. Most previous work has examined either global network dynamics or local neural patterns separately, leaving questions about how these scales relate to each other open.
- Third, we study whether transitions between anti-correlated states show systematic relationships with stimulus features, including auditory and visual changes, as well as event boundaries. Previous work has typically investigated only a single stimulus dimension (***Baldassano et al., 2017***; ***Geerligs et al., 2022***) and has not investigated how these associations are tied across spatial scales.

To address these aims, we used data from 15 participants who watched a slightly edited version of the movie “Forrest Gump”, while their brain activity was recorded with 3T fMRI (StudyForrest dataset; ***Hanke et al., 2016***). These data were hyperaligned and group-averaged, which allows us to examine stimulus-driven neural dynamics while minimizing idiosyncratic individual differences. To investigate the consistency of our findings, we also replicated our analyses in two different 3T fMRI datasets where participants were either watching an 8 minute movie (CamCAN data, 285 participants, ***Shafto et al., 2014***; ***Taylor et al., 2017***) or listening to three audio narratives of 10 minutes each (Narrattention, 54 participants). This allowed us to determine whether these local and global anti-correlated dynamics represent a general organizational principle of naturalistic neural processing.

## Results

### Two anti-correlated states dominate naturalistic continuous fMRI activity

#### Global

To investigate the anti-correlated nature of neural states at the scale of the whole brain, we first take a closer look at the correlational structure of continuous whole-brain fMRI activity. To this end, we compute the average brain activity across participants in a movie-watching fMRI dataset (StudyForrest). This provides us with the brain-activity that is evoked by the stimulus in a consistent way across participants. Next, we create a whole-brain (region of interest (ROI) by time) timeseries by averaging BOLD activity across the voxels within each of the 400 parcels of the Schaefer Atlas (***Schaefer et al., 2017***) (see Figure 1a). To investigate whether the brain activity visits repeating neural states, we can compute the correlations between neural patterns at each timepoint, i.e. a time-point by timepoint correlation matrix (see Figure 1b). One recognizable feature of this time by time correlation matrix is that the majority of neural activity patterns share a common correlation structure to other neural activity patterns. This structure appears as transitions from highly positive to highly negative correlations (or vice versa) with relatively stable periods in between. It is present for almost all timepoint correlations, suggesting that two main opposing patterns are expressed at different timepoints. This implies that brain activity alternates between two anti-correlated states in movie watching fMRI data.

The time by time correlation matrix shows the direct relationships between neural activity patterns at each timepoint. To understand how these relationships themselves are correlated, we can compute a ‘meta’ temporal-correlation matrix which shows the degree to which the rows of our temporal correlation matrix are similar to each other (see Figure 1c). Each element in this matrix now tells us how similar neural patterns at two timepoints are in terms of their relationship to all other timepoints. When two timepoints are strongly correlated in the meta-correlation matrix, the neural patterns at those timepoints share a similar ‘fingerprint’ of how they relate to patterns at other times. In other words, even if their specific neural activity patterns are different, they play similar roles in the overall temporal correlation structure of our signal.

This ‘meta’ correlation matrix shows a great deal of structure: timepoints have long-range correlation structures with one another and the majority of timepoints are either strongly correlated or strongly anti-correlated to each other. Here, we only plotted a small portion of all timepoints in the dataset however we observed the same structure across different segments of the timeseries. To test whether these observations hold across the whole dataset, we applied a statistical test of stationarity. We used the Augmented Dickey-Fuller Test (***MacKinnon, 1994***) to test the stability of the temporal correlation structure in the data and see that the correlation structure is indeed stationary (p < 0.001). This indicates that the observed long-range temporal correlations represent stable, persistent patterns rather than non-stationary trends or random walks. Therefore we can conclude that the anti-correlational structure is long range and not localized to a particular time period: it is a general property of continuous whole-brain BOLD activity in naturalistic settings.

Looking at the ‘meta’ correlation matrix at Figure 1c, we can see that neural patterns at most timepoints are highly positively or negatively correlated with patterns at other timepoints. For timepoints that show this behaviour, we expect the correlation values to show a bimodal distribution with many positive and negative correlations to other timepoints and fewer correlations around zero. To investigate if this is the case, we applied a dip-test for bimodality to the meta temporal correlation values of each timepoint. We found evidence for a bimodal distribution in the majority (63%) of timepoints (with 82% of all significant p-values being smaller than 0.0001), reflecting that neural activity has an underlying multivariate signal which flips between two strongly anti-correlated states. We consider all timepoints which lie on the positive and negative parts of the bimodal distribution as belonging to distinct anti-correlated states (shown as the brown and teal distribution in Figure 1d). We refer to the remaining timepoints (37%) as the unimodal state, as they do not significantly participate in the positive/negative correlation pattern (shown as the green histogram).

To understand the temporal dynamics of these three states (positive, negative, and unimodal), we label all fMRI timepoints based on our analyses so far and create a timeline of states (Figure 1e). The transition matrix of this timeline (Figure 1d, right), shows that there are few transitions from one bimodal state into the other bimodal state. Rather, transitions tend to occur via the unimodal state. The unimodal state, on the other hand, has equal transition probabilities to the two ends of the bimodal states. This suggests that the brain spends most of its time in one of the two anti-correlated states, with some non-repeating unique states that mediate the transitions in between. Overall, the brain spends roughly equal time in all three states (Figure 1d, middle). Figure 1e shows the progression of the whole-brain activity over time in relation to the progression of states that is shown below.

As we will show in the following sections, the cortical maps of these two bimodal states reveal a clear separation of default mode areas and task positive areas, also see Appendix 1. Therefore we label the three states we have just described as: Task Positive State (TPS), Default Mode State (DMS), and Unimodal State.

#### Local

So far, we have investigated whole brain activity patterns based on ROI averaged whole-brain signals. To see if our observations also hold for local activity patterns, we performed the same analyses for voxel-wise activity patterns within ROIs. We observed the same anti-correlated structure within ROIs as we found on the global scale (see Figure2c and Appendix2) and each of these exhibited stationary correlation structures (all p-values < 0.005). Similar to the whole-brain results, we found a strong bimodal distribution of states in ROI timeseries. Figure 1f shows the proportion of the three states for all ROIs. Compared to the global whole-brain signal, voxel-wise local activity spends less time in unimodal states for most ROIs (Figure 1f). When we compare the two bimodal states, we observe that there is always one state that has higher average brain activity than its anti-correlational counterpart (both for the global and the local data).

To test for significant mean BOLD activity differences local anti-correlated states, we performed Welch’s t-tests (assuming unequal variances) comparing the mean activities of timepoints that are labelled as the two different states across all ROIs. The results show highly significant differences in mean activity across the two states for nearly all ROIs (97,5%, see Figure 1g). As local anti-correlated states correspond to high and low BOLD activity timepoints, we labelled accordingly as up, down, and unimodal states (as opposed to TPS, DMS, and unimodal at the global scale). It should be noted that there are also a few ROIs which spend the majority of their time in the unimodal states. Specifically the orbitofrontal cortex and the temporal poles do not have any bimodal states and only operate in unimodal states. These two ROIs are specifically known to have unreliable BOLD signals due to signal loss, of which this may be a reflection.

In summary, we have shown that naturalistic brain activity can be grouped, across spatial scales, into three broad categories: the either ends of a repeating anti-correlational template pattern and non-repeating transitions in between. In the remainder of this manuscript we focus on the anti-correlated states.

### Creating Template Patterns of The Anti-Correlated Signal

#### Global Anti-Correlated States

Turning our attention to the bimodal states, we next investigate the core spatial pattern of activity that contributes to the switch between these anti-correlated states. To do so, we fit a linear model to the timepoints corresponding to the extreme ends of this bimodal distribution of states. This extracts a ‘flipping’ template pattern which describes the multivariate pattern of neural activity which ‘flips’ in an anti-correlated way and transitions between these two states. In the remainder of the manuscript we use this template pattern to investigate the spatial and temporal properties of local ‘up’ and ‘down’ states as well as global default mode and task positive states (DMS and TPS).

By fitting our neural data at each timepoint to these template patterns, we can see that our template measures the global alignment of timepoints with the default mode state when negatively loaded (i.e., having a negative weight in the template) and with the task positive state when positively loaded (Figure 2b). Similarly at a local level, the template magnitude aligns with up and down states.

**Figure 2.**
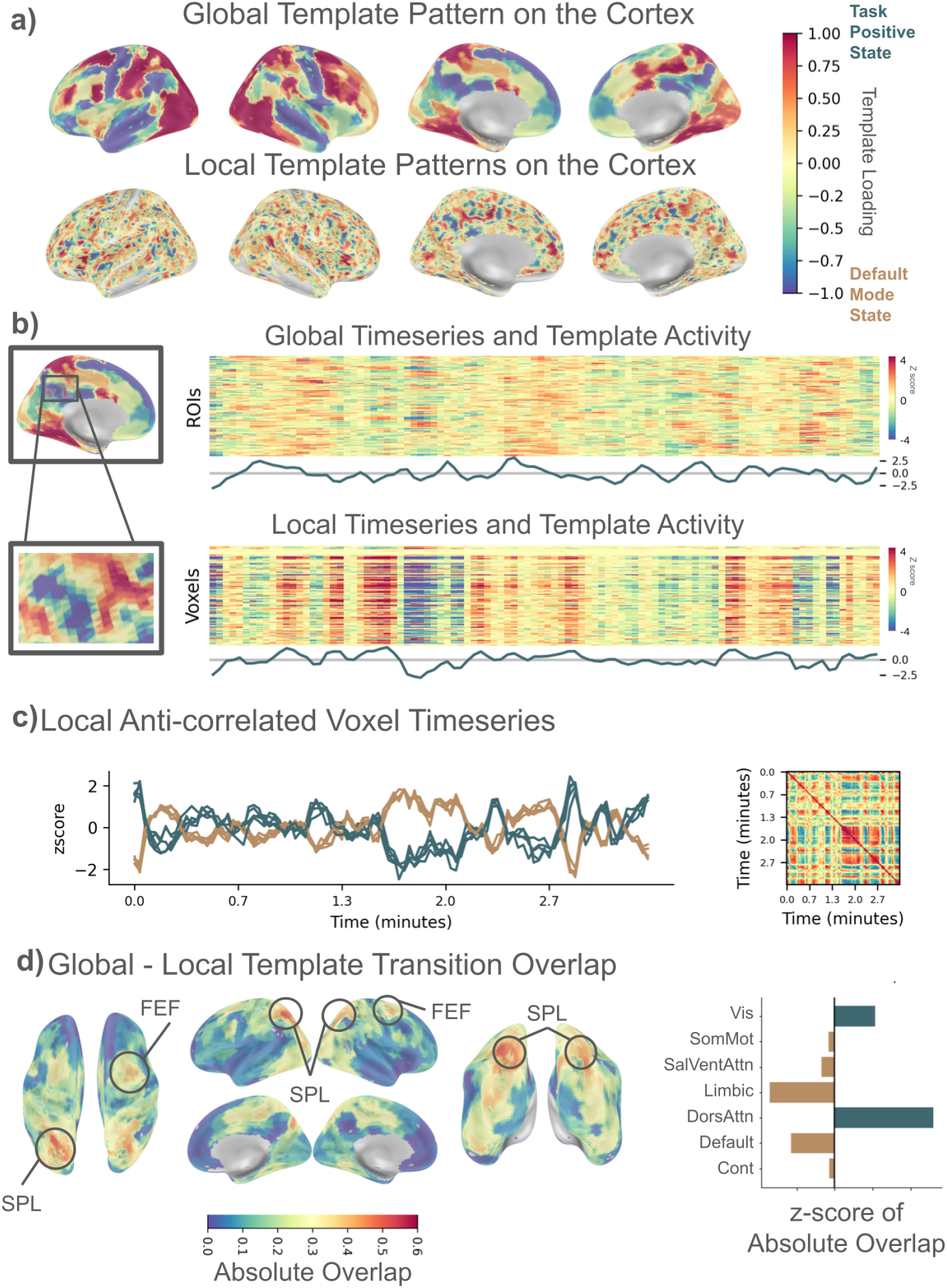
Anti-Correlated signals across spatial scales are related. (a) Global and local anti-correlated patterns mapped on the whole cortex. Top: Global anti-correlated pattern which differentiates the Default Mode State (blue) and Task Positive State (red) visualised on the cortex. Bottom: Local anti-correlated patches of voxels visualised on the cortex. (b) Top: Global whole-brain (ROI averaged multivariate timeseries), and Bottom: an example ROI’s local (within ROI multivariate voxel timeseries) timeseries is plotted above the anti-correlated fluctuations of the template shown in line plots. (c) Left: Ten example voxel timeseries from the left parietal inferior lobule which were selected at the extreme ends of the template patterns to illustrate the presence of locally anti-correlated voxels, and Right: time by time correlation matrix of left parietal inferior lobule. (d) Global - Local anti-correlated state switches are most strongly aligned in the Dorsal Attention Network (DAN) areas, specifically FEF and SPL

To evaluate how well the global template represents the underlying neural dynamics across brain regions, we quantify the reconstruction quality by looking at the proportion of variance explained. The global template preserved 26% of the variance in the original ROI-averaged data, indicating moderate reconstruction quality. The global mean activity on the other hand preserved only 17% of the variance. This suggests the template captures key shared patterns across ROIs while individual ROI-specific variance remains largely unexplained.

The global template pattern mapped onto the cortex is shown in Figure 2a, top row (an annotated version of this map can be found in Appendix 1a). Areas that have a negative loading in this template pattern are mostly active in the default mode state, while areas that have a positive loading are mostly active in the task positive state. Specifically, the areas most active in the task positive state are the visual cortex, occipito-temporal and occipito-parietal cortex, superior parietal lobules, postcentral gyri, frontal eye fields, parietal operculi, parts of the frontal and parietal medial cortex, parts of the precuneus, and parts of the intraparietal sulcus. Together, these cover large parts of what has been termed the task positive network (TPN). Indeed, when we investigate the template loading within each of the major functional brain networks defined by ***Thomas Yeo et al. (2011***), we see contributions of the dorsal attention network (DAN) and less so the visual network to the task positive state (see Appendix 1b). The areas most active in the default mode state on the other hand consists of the precuneus, angular gyrus, large parts of the superior and middle temporal cortex, large parts of the somatomotor areas, frontal operculi, insula, parts of the prefrontal cortex and limbic areas. These regions cover large parts of the default mode network but also include typically unrelated areas such as the somatomotor system (see Appendix 1b). Overall, 69% of all ROIs are associated with the TPS, and 30% with the DMS.

These results are in line with previous literature on global brain states (***Yamashita et al., 2021***; ***Song et al., 2023***). Particularly, the first cortical connectivity gradient described in ***Margulies et al. (2016***), as well as the quasi-periodic patterns (QPPs; ***Thompson et al., 2014***; ***Yousefi and Keilholz, 2021***; ***Majeed et al., 2011***) both demonstrate this anti-correlational global transition between the two states in resting state fMRI. These two states show strong spatial correspondence to the two ends of the global template pattern that we observe. Indeed, when we computed QPPs in our dataset, we observed that these were highly similar to the template patterns we describe here (see Appendix 6).

#### Local Anti-Correlated states

Next, to understand the spatial distributions of local anti-correlated activity, we investigate the same template patterns within local ROIs (for an example, see Figure 2b, bottom row). To do so, we plot the voxel-wise ROI template patterns (template loading coefficients) for all ROIs, covering the whole cortex. This map uncovers a mesoscale functional organisation of the entire cortical sheet (see Figure 2a bottom). Patches of voxels with positive template loadings are interspersed with patches of voxels that have negative template loadings. This means that when we look at local activity, nearby voxels can respond to the same stimulus input in an anti-correlated way with respect to their neighbours (for examples, see Figure2c and Appendix2). Within ROIs, an average of 73% of voxels have positive loadings (with a standard deviation of 14%). Around 27% of voxels show an opposite response, showing that they respond in an anti-correlated way compared to the majority of voxels in an ROI.

Note that the spatial distribution of local template loadings looks different from the global template for two reasons. First, the resolution of these patterns differs. Global patterns are measured based on ROI-averaged data, while local patterns are computed voxel-wise within individual ROIs. Second, and more importantly, the global and local templates use different references. The global template considers a single template that can explain whole brain changes in activity across a major anti-correlational axis. Locally, a template is constructed independently for each ROI to capture its own local axis of anti-correlation in voxel-wise activity.

To evaluate how well the local templates represent the underlying neural dynamics within brain regions, we quantified the reconstruction quality using the proportion of variance explained metric. The local templates preserved on average 44% of the variance in the original within-ROI data with a standard deviation of 20% across ROIs. The local mean activity on the other hand preserved only on average 27% of the variance in the original within-ROI data with a standard deviation of 18% across ROIs.

When we compute a global template based on voxel-wise activity, we can also see such a local patterning of cortical alignment with the extracted template, but with less contrast than the local referencing shown here (see Appendix 2). To ensure that the local template patterns are not a product of hyperalignment, we also ran the same analysis on non-hyperaligned data (see Appendix 2). We see that the fine mesoscale organisation seen in the local template patterns with hyperaligned data is still present in non-hyperaligned data, but the structure is coarser with bigger anti-correlated patches. This indicates that our results are not a by-product of hyperalignment, but hyperalignment allows us to observe finer spatial details due to improved alignment across participants.

#### Anti-correlation across spatial scales

The results so far show that as the whole brain switches from one anti-correlated state to another, local areas also go through switches between anti-correlated states of their own, highlighting a nested organization in cortical activity. To characterise the relationship between the global and the local anti-correlational activity in more detail, we calculated the absolute overlap (***Geerligs et al., 2022***) of the transition moments between global and local states. The absolute overlap metric (OA) measures overlap between two different ‘boundary’ annotations by using the minimum number of boundaries of either type as its reference, and normalizing the difference between this minimum and the expected overlap. Here a transition moment is defined as a moment when the template timeseries crosses zero. Figure 2c shows the cortical map of the absolute overlap ratio between the global and local transitions. We can see that the superior parietal lobe (SPL) and the frontal eye fields (FEF) show the greatest overlap between their local ROI state switches and the global state switches. Figure 2c shows that, overall, the Dorsal Attention Network (DAN) areas show the highest synchrony with the global states. These suggests that the transitions in global dynamics could be the result of localized interactions, mainly focused in the DAN.

### Anti-correlation in relation to stimuli

To understand the relationship between the anti-correlated states and the stimuli better, we measured which of the two anti-correlated states was more prevalent at the whole-brain scale and in local ROIs at moments of stimulus change. It is important to note that all of these analyses are done on group averaged movie-viewing data. This means that our brain signals are driven by shared responses to stimuli, as endogenous ongoing activity is averaged out. To investigate neural states at moments of stimulus change, we examined the neural data for three different stimulus annotation types: transitions between shots (i.e. transitions from one continuous piece of film to another) to focus on the visual modality, changes in the Mel-Frequency Cepstral Coefficients (MFCC) of the audio to focus on the auditory modality, and behaviourally annotated event boundaries to focus on multimodal, abstract features. To account for the delay in the HRF, each of the stimulus onsets were adjusted by 4.5 seconds.

For both whole-brain and local analyses, we used a permutation-based statistical test. We calculated the overlap between detected neural states and stimulus boundaries, then compared this to null distributions generated through 10,000 permutations. These permutations preserved the temporal structure of the stimuli by maintaining inter-boundary durations while shuffling their order. We examined state transitions within temporal windows of ±10 timepoints (TRs) for whole-brain and ±3 timepoints for regional analyses, applying false discovery rate (FDR) correction for multiple comparisons.

As can be seen in Figure 3 b, d, and f, all stimulus feature types triggered significant global shifts toward the TPS, though with varying magnitudes and durations. Shot transitions showed the strongest effect, with a 40% increase in TPS probability compared to baseline (p<0.001), and this elevated probability persisted for four TRs (8 seconds). Event boundaries produced a more moderate effect with a 21% increase in TPS probability (p<0.01), lasting for (6 seconds) three TRs. MFCC changes showed the smallest but still significant effect with an 18% increase (p<0.05) lasting two TRs (4 seconds). This pattern is also reflected in the local up state prevalences, with shots leading to a longer wider-ranging set of areas to be in the up state which is also sustained for longer, which then likely translate as more prominent global state changes.

**Figure 3.**
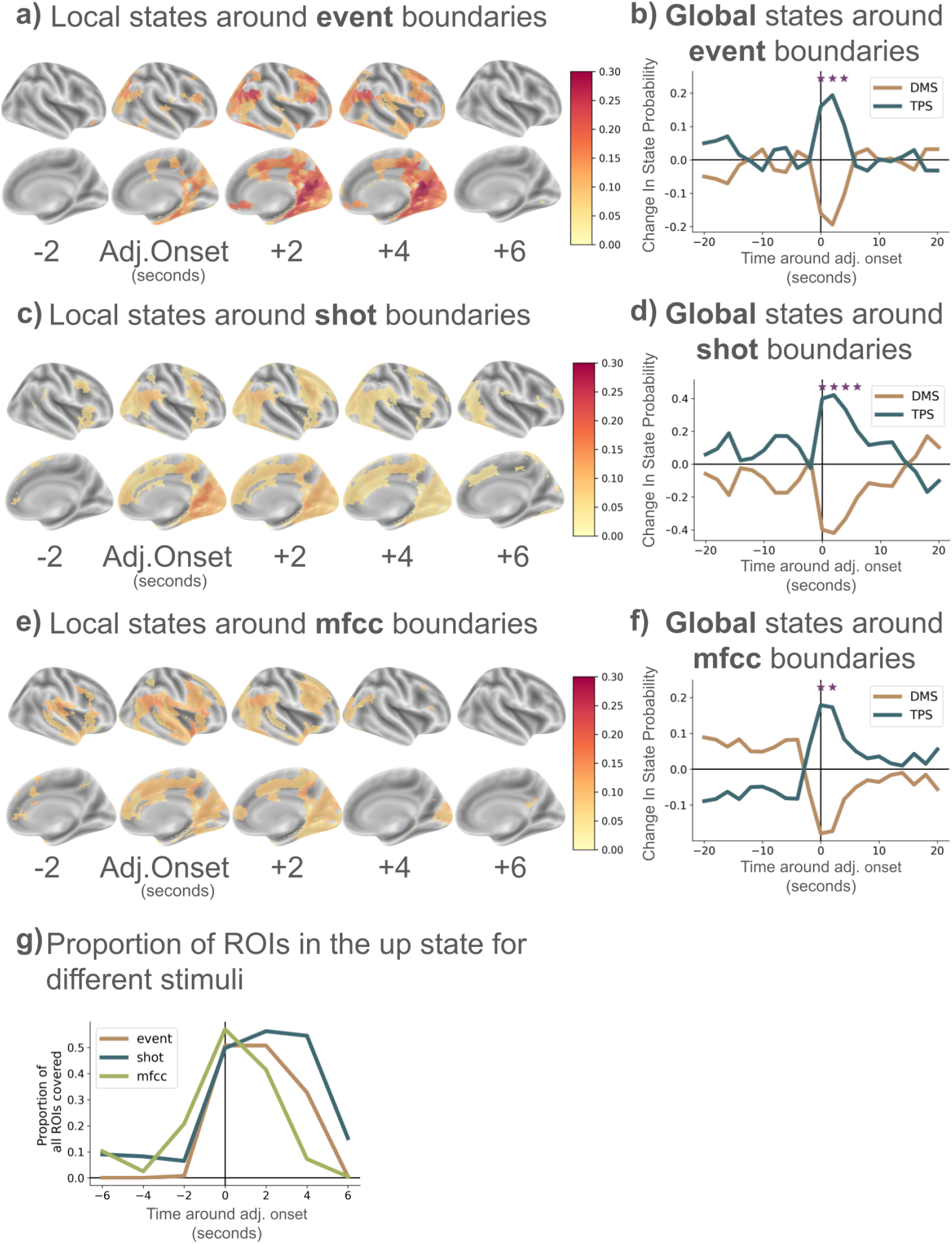
Stimuli boundaries are followed by a global transition to the TPS, and coincide with transitions to localized up states. (a) Increase in probability of being in the up state spreads across the cortex after event boundaries. (b) Probability of being in the TPS increases after event boundaries. (c) Increase in probability of being in the up state compared to baseline spreads across the cortex after shot changes. (d) Probability of being in the TPS increases after shot boundaries. (e) Increase in probability of being in the up state spreads across the cortex after MFCC changes. (f) Probability of being in the TPS increases after MFCC boundaries. (g) Stimuli changes lead to a cascade of local up state transitions across the cortex and then dissipate. Results are visualized as state probability changes around boundaries, with significant timepoints (p<0.05 after FDR correction) marked with asterisks

Figure 3 a, c, and e demonstrate modality-specific effects of stimuli on brain activity. MFCC boundaries predominantly trigger up states across the middle and superior temporal cortex extending toward the occipital cortex and inferior frontal areas bilaterally (Figure 3e), while also showing statistically significant but smaller effects across parts of the occipital cortex, in medial parietal areas and the dorsolateral prefrontal cortex. Shot boundaries (Figure 3 c) trigger up states across the entire occipital cortex, but also extending into lateral and medial parietal regions as well as medial prefrontal cortex and inferior frontal cortex, with minimal significant effects in temporal areas related to audition. Event boundaries (Figure 3 a), defined by subjective human ratings rather than being strictly perceptual features, are associated with a stronger probability of being to the up-state in many of the areas that are also driven by shot-changes. However, event boundaries are related to stronger probabilities of being in the up-state. It should be noted that in naturalistic settings such as movie-watching there is considerable overlap between onsets of different stimuli types. For example 60% of event boundaries overlapped with shot boundaries and 26% with MFCC boundaries, 33% of MFCC boundaries overlapped with shot boundaries while 15% of shot boundaries overlapped with MFCC boundaries. This makes it difficult to fully separate effects of visual changes, auditory changes or event boundaries (***Oetringer et al., 2025***). Nevertheless, the responses to MFCC and shot boundaries show clear evidence for a modality specific response.

Figure 3g shows how the proportion of all ROIs which are statistically significantly more in an up state change in time for different stimuli boundary types, and demonstrates a clear temporal progression in stimulus-evoked responses. For all stimuli types, approximately 50% of all ROIs were significantly more likely to have transitioned to the up state immediately following the HRF-adjusted stimulus onset. The effect dissipated within two to three TRs as regions gradually returned to their baseline up state probabilities, with shot changes maintaining elevated activity longer.

These findings establish a direct link between the anti-correlated global and local states and the processing of external stimuli. The predominance of the “up” states across wide sensory and trans-modal areas after stimuli boundaries suggests that when the relevant aspects of the environment change, processing demands in corresponding areas go up. Similarly, the predominant activation of visual areas in response to shot changes and the predominant activation of the auditory areas with MFCC changes reinforces the idea that local state dynamics are related to modality-specific processing demands. Eventually, noticeable stimuli changes might trigger an up state transition in a large-enough proportion of the cortex that they often lead to a global brain state change. Alternatively, global brain state changes might trigger up or down state transitions in local cortical areas. Future work should investigate the causal structure between global and neural states. Together, these results extend the evidence for a nested spatio-temporal structure in naturalistic neural activity, where local processing demands can cascade into global state transitions when stimulus changes are sufficiently salient, and/or global states might influence local states; likely both depending on the context.

### Replication and Summary of Findings Across Diverse Datasets

To test the generalizability of our methods and results beyond a single dataset, we extended our analyses to two additional naturalistic fMRI datasets. From the CamCAN dataset, we use the data of 265 subjects who viewed an 8-minute version of Alfred Hitchcock’s black-and-white television narrative “Bang! You’re Dead” (***Shafto et al., 2014***). The other dataset we used was collected in the context of another ongoing research project (Narrattention data). Here, we use data of 54 participants who listened to 3 Dutch stories (audiobooks, 18 participants per story) which were approximately 10 minutes each.

We replicated the key findings of this paper with these different datasets, albeit with some differences, as described below:

1. **Anti-correlated patterns are a dominant feature of neural activity during movie watching and story listening**. Across all datasets, our results show that continuous brain activity organizes into three broad categories: two extremes of a repeating anti-correlational pattern, and non-repeating transitions between them, as demonstrated across all datasets in Figure 4 a & b. Globally, these patterns manifest as whole-brain states that correspond to the DMN and TPN configurations, which alternate in a semi-regular fashion with transition states mediating between them. The consistency of this pattern across datasets indicates that this organization represents a fundamental property of brain function during naturalistic conditions rather than a dataset/task-specific phenomenon. Locally, individual ROIs exhibit similar anti-correlated activity patterns at a finer spatial scale across all datasets, with neighbouring voxels showing opposite response patterns. The replication across datasets with different stimulus types (audiovisual and purely auditory) indicates that anti-correlated neural dynamics represent a domain-general organizational principle rather than a modality or task -specific phenomenon.

**Figure 4.**
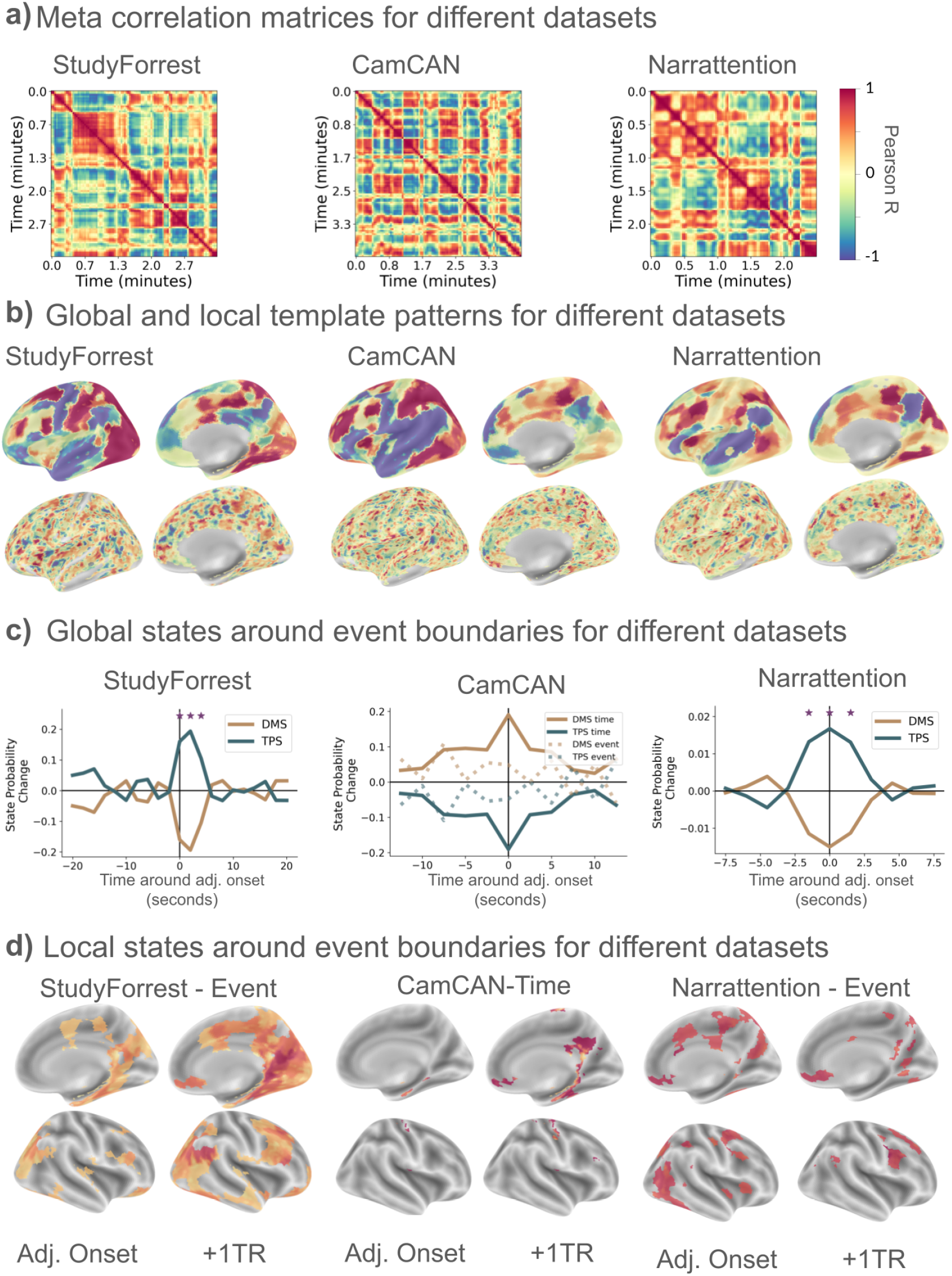
Results replicate across different datasets. (a) StudyForrest, CamCAN, and Narrattention datasets all display anti-correlated meta correlation matrices. (b) Global anti-correlated states extracted from the StudyForrest, CamCAN, and Narrattention datasets are consistent. (c) Change in probability of being in the TPS increases compared to baseline after event boundaries for both StudyForrest and Narrattention. Significant timepoints are marked with asterisks (p<0.05 after FDR correction) (d) Increase in probability of being in the up state spreads across the cortex after event boundaries for both StudyForrest and Narrattention.
2. **Anti-correlated neural states at global and local scales are related to one another**. The overlap between boundaries at the local and global scales also replicates across datasets, although the precise areas that show most overlap differ (see 5). Specifically while the DAN areas show the highest overlap with global state changes for StudyForrest and Narrattention, for CamCAN it was the ventral attention network (VAN). The difference between these dataset suggests that the coupling between global and local neural states may be partially determined by the stimulus characteristics.
3. **Transitions between anti-correlated states show systematic relationships with stimulus features**. In the Narrattention dataset, we observe an increase in the occurrence of the TPS state at event boundaries, similar to what we found in the StudyForrest data (see figures 4c). It is worth noting that the Narrattention dataset contains neural activity elicited solely by auditory stimuli. This suggests that the association between TPS and event boundaries cannot be explained only by shot changes or other visual changes. In contrast, we did not observe a significant association between event boundaries and global state probability in the CamCAN data, which might be the result of the short duration of the movie stimulus. When we instead looked at a more objective annotation of changes in time in the stimulus (frames after a cut is temporally discontinuous with the frame preceding the cut) we observed a more prominent response, but not statistically significantly, and with an increase of DMS probability. This could suggest that differences in the properties of the movie could determine which state is more prevalent at a boundary.

At the local scale, we see increases in the occurrences of up-states in the Narrattention dataset, that overlaps with many of the areas we observed in the StudyForrest data (see 4d and 1)). However, the effect is weaker than for StudyForrest, possibly because of the shorter stimulus duration. For CamCAN, we see the same, but here with only a few significant voxels (1). When we again investigate objective annotation of changes in time in the stimulus, we find that areas in the precuneus tend to be in the up-state just after a time-shift (see 4d). The increased likelihood of either of the global anti-correlated states around event boundaries, independent of sensory modality, shows that global brain states are an important neural mechanism in event cognition. This suggests that the mechanisms underlying these anti-correlated states likely serve fundamental cognitive functions related to information processing, attention, and event segmentation.

Our analysis pipeline is available as a Jupyter notebook, and we encourage readers to replicate our findings using their own datasets, as the approach is expected to generalize to alternative data with comparable results.

## Discussion

Our investigation of naturalistic brain activity during movie viewing has revealed a fundamental organizational principle that spans spatial scales: the oscillation between two anti-correlated neural states, mediated by a transitional state. This tripartite structure, evident at both global network and local circuit levels, suggests a nested hierarchy of brain dynamics that balances internally and externally focused cognition. The robustness of these findings across multiple independent datasets, including those with different sensory modalities, indicates that this phenomenon represents a domain-general feature of neural processing rather than an effect specific to a particular stimulus. In the following discussion, we first situate these findings within the extensive literatures on the antagonistic relationship between the Default Mode Network and task-positive systems, and quasi-periodic patterns (QPPs). We examine how these anti-correlated states implement a fundamental operating rhythm for naturalistic cognition, with each state serving complementary cognitive functions. We then explore how these dynamics manifest at the local level. Finally, we consider the theoretical implications of anti-correlated neural dynamics as a scale-invariant organizational principle and draw connections to computational frameworks from artificial neural networks that implement similar principles. Throughout, we propose that anti-correlated dynamics represent a convergent solution to the fundamental challenge of maintaining coherent internal representations while remaining responsive to meaningful changes in the environment. This mechanism may underlie the brain’s ability to segment continuous experience into meaningful events while preserving narrative continuity.

### Global Anti-correlated Activity

Our findings of whole-brain anti-correlation that is driven by states in which either the DMN or the TPN is more active, aligns with early work that characterized the antagonistic relationship between these networks (***Fox et al., 2005***; ***Raichle et al., 2001***; ***Greicius et al., 2002***). Our results suggest that this antagonism manifests as a fundamental organizational principle of brain activity also during naturalistic viewing, where discrete shifts between these anti-correlated states occur regularly.

#### Default Mode State

The DMN was initially discovered as a set of brain regions that consistently show deactivation during externally focused tasks (***Raichle et al., 2001***; ***Greicius et al., 2002***). It is comprised of the medial prefrontal cortex, posterior cingulate cortex, angular gyrus, and medial temporal lobe structures. Recently, it has been increasingly recognized as playing a central role in internal mental processes such as self-reference, social cognition, episodic memory, and language comprehension (***Menon, 2023***), in line with its position at the top of a cortical gradient supporting abstract processing (***Margulies et al., 2016***).

In line with this, converging evidence suggests that key areas of the DMN serve as a cognitive architecture that actively constructs and updates situation models while representing events at multiple temporal scales (***Nguyen et al., 2019***; ***Baldassano et al., 2017, 2018***; ***Simony et al., 2016***). Together, these findings suggest that the DMN supports abstract processing, where it bridges stored knowledge with ongoing experience and enables knowledge transfer across different contexts (***Binder et al., 2009***; ***Spreng et al., 2009***). We observed that the DMS tended to be less present at moments where there were event boundaries or large changes in the visual or auditory components of the stimulus, suggesting that this state may be more prevalent during stable periods where little change occurs in the external environment. However, in the CamCAN data we observed that the DMS was more likely to occur after a shift in time in the narrative and previous work also observed an increase in the DMN at event boundaries (***Song et al., 2023***). We may speculate that event model updating can drive the brain to the DMS, however, when this is accompanied by many changes in sensory input the TPS may be dominant instead.

#### Task Positive State

The Task Positive State (TPS) observed in our study represents a configuration of brain activity characterized by heightened engagement of sensory, attention, and executive control networks (***Fox et al., 2005***; ***Raichle et al., 2001***; ***Weigard et al., 2024***; ***Cheng et al., 2020***; ***Barber et al., 2016***; ***Hammer et al., 2024***; ***Leech et al., 2025***; ***Kucyi et al., 2020***). This state aligns with what has been traditionally termed the Task Positive Network (TPN), including regions such as the dorsal attention network (DAN), control network (CN), and visual areas. Our findings demonstrate that the TPS becomes more dominant immediately following perceptual changes in naturalistic stimuli (e.g., visual scene changes, auditory transitions) and event boundaries, suggesting its role in processing and responding to salient changes in the environment.

The functional profile of the TPS corresponds well with existing literature on externally-directed cognition. Multiple studies have shown that these networks are preferentially engaged during tasks requiring focused attention, working memory, and cognitive control (***Fox et al., 2005***; ***Raichle et al., 2001***; ***Weigard et al., 2024***; ***Cheng et al., 2020***; ***Barber et al., 2016***; ***Hammer et al., 2024***; ***Leech et al., 2025***; ***Kucyi et al., 2020***). Importantly, our results extend these findings by demonstrating the dynamic engagement of the TPS during naturalistic viewing without explicit task demands, suggesting it represents a fundamental mode of brain function engaged automatically in response to meaningful environmental changes.

#### Interaction between the two

The anti-correlated relationship between the Default Mode Network and Task Positive Network has been a foundational observation in functional neuroimaging since it was first identified by (***Fox et al., 2005***). This intrinsic anti-correlation has been consistently observed across numerous studies in resting state (***Fox et al., 2005***; ***Fransson, 2005***), task-based paradigms (***Kelly et al., 2008***), and has been linked to behavioural performance (***Hampson et al., 2010***; ***Keller et al., 2015***). What our study contributes is evidence that this well-established anti-correlation manifests as a dynamic alternation between discrete global brain states during naturalistic stimulation, with systematic transitions triggered by environmental changes. This does not imply that these systems are necessarily always anti-correlated as we found that around 34% of the time, the brain was in a unimodal state, in line with previous research has shown that these networks can activate together depending on the task context (***Dixon et al., 2017***; ***Spreng et al., 2010***).

#### QPPs

The ongoing shifts between states with activity dominated by either the DMN or the TPN is very similar to QPPs that have been described in resting state and task settings. Initially identified in mice by ***Majeed et al. (2011***), QPPs were described as recurring cycles of activation and deactivation between the DMN and TPN, repeating roughly once in twenty seconds. These patterns were later confirmed in humans and linked to large-scale network dynamics that alternate between internally and externally directed cognitive processes (***Thompson et al., 2014***; ***Yousefi and Keilholz, 2021***). The physiological underpinnings of QPPs, as highlighted by ***Thompson et al. (2014***), linked QPPs observed in fMRI to infraslow local field potentials (LFPs), hinting at a shared mechanism between electrophysiological oscillations and BOLD-based spatiotemporal patterns. Moreover, recent studies such as those by ***Yousefi and Keilholz (2021***) and ***Bolt et al. (2022***) have shown how QPPs propagate along macroscale gradients and represent canonical spatiotemporal patterns coordinating functional connectivity across the brain. By replicating these findings in naturalistic settings, while isolating stimulus-driven activity, we extend their relevance, showing that QPP-like dynamics underlie the brain’s adaptability to naturalistic environmental stimuli. These results validate QPP-like dynamics as a universal organizing principle of brain function, bridging insights across methodologies, cognitive states, and spatial scales.

The methodological advances in our work offer key improvements to the traditional analysis of QPPs. While prior studies relied on template-matching algorithms with fixed oscillatory durations (***Majeed et al., 2011***; ***Thompson et al., 2014***; ***Yousefi and Keilholz, 2021***), we allowed oscillatory dynamics to emerge naturally from the data, unconstrained by predefined parameters. This approach provides a more precise characterization of the temporal evolution of global states, capturing variability in the duration and timing of state transitions. Furthermore, we demonstrated that the same analytical framework used to detect global QPPs can be extended to local cortical regions, uncovering a nested hierarchy of neural state dynamics. Specifically, our results link local voxel-level anti-correlations to the broader global patterns of DMN-TPN alternations, highlighting how local and global dynamics integrate. This advancement bridges a gap in the literature, which has predominantly examined either global or local dynamics in isolation, and reveals the hierarchical organization of QPP-like activity across scales.

Our results indicate that the DMS-TPS alternation appears to implement a fundamental operating rhythm for naturalistic cognition. The brain cycles between these two complementary processing modes: periods of TPS dominance for active processing of new perceptual information, and periods of DMS dominance that may support integration and maintenance of contextual information. The temporal structure of these transitions, with salient perceptual changes triggering a temporary shift toward TPS dominance, followed by a return to DMS dominance, suggests a general processing loop for naturalistic cognition.

### Local Anti-correlated Activity

Local anti-correlated activity have also been studied indirectly in the context of neural states. Literature shows that under naturalistic conditions local brain activity activity transitions from one temporarily stable activity pattern to another, with fast transitions in between ((***Baldassano et al., 2017***; ***Geerligs et al., 2022***; ***Oetringer et al., 2025***). In other words, it progresses as a series of stable states. ***Geerligs et al. (2022***) demonstrates that neural states are organized in a temporal cortical hierarchy, with short states in primary sensory regions, and long states in lateral and medial prefrontal cortex. Here, we show that these states are mainly driven by two distinct populations of anti-correlated voxels, mirroring the anti-correlated networks we see at the global scale. Projected on the whole cortex, these populations reveal a mesoscale functional organisation consisting of patches of anti-correlated voxels. The distinct anti-correlation between these voxel patches/clusters might relate to local inhibitory connections underlying this phenomenon (***Cardin et al., 2009***; ***Sohal et al., 2009***).

Our analyses on local state transitions in response to different types of stimulus changes revealed a clear pattern of modality-specific responses, with visual regions showing more prominent transitioning to an “up state” following visual changes (shot boundaries) and auditory and language-related regions showing similar transitions following auditory changes (MFCC boundaries).

The selective activation of sensory areas in response to changes in the relevant sensory information reflects the well-established organizational principle of functional specialization in the brain. Here we show that these modality-specific responses are implemented through transitions between anti-correlated states within local neural populations.

### Interaction between global and local

Perhaps most significant is our finding that transitions between anti-correlated states are often synchronized across spatial scales, particularly in regions like the frontal eye fields (FEF) and superior parietal lobule (SPL), which are part of the Dorsal Attention Network (DAN). This network, traditionally associated with top-down directed attention (***Corbetta and Shulman, 2002***; ***Vossel et al., 2013***), may serve as a key interface between local perceptual processing and global state dynamics. Both the StudyForrest and Narrattention dataset showed the highest overlap between local and global neural state changes in DAN areas, suggesting that these areas may detect salient changes in the environment and initiate cascading transitions across spatial scales. This aligns with the DAN’s established role in orienting attention to behaviourally relevant stimuli and maintaining goal-directed cognitive states (***Vossel et al., 2013***). The DAN’s position at the interface between sensory and higher-order cognitive systems makes it ideally situated to propagate state transitions from local sensory regions to global network configurations (***Margulies et al., 2016***; ***Sepulcre et al., 2012***; ***Corbetta and Shulman, 2002***; ***Vossel et al., 2013***; ***Shulman et al., 2009***). However, these results were not replicated in the CamCAN dataset, suggesting that the overlap between global and local changes may also depend on the characteristics of the stimulus.

This synchronization between global and local state changes suggests a mechanism for integrating local and global processing: when global state transitions occur, they may cascade to or emerge from coordinated transitions in local regions, allowing for coherent updating of both perceptual details and overarching narrative frameworks. This integration across scales may be bidirectional. Bottom-up propagation would allow locally detected perceptual changes (e.g., in sensory regions) to trigger global state transitions when sufficiently salient, enabling narrative updating in response to significant perceptual shifts. Top-down propagation would permit global state changes to synchronize processing across distributed local regions, ensuring that contextual updating occurs in a coordinated fashion throughout the brain. The DAN regions, showing the strongest synchronization between local and global transitions, are notably involved in attention control and multimodal integration (***Margulies et al., 2016***; ***Sepulcre et al., 2012***; ***Corbetta and Shulman, 2002***; ***Vossel et al., 2013***; ***Shulman et al., 2009***), positioning them ideally to serve as interfaces between local perceptual processing and global narrative frameworks. These regions may detect mismatches between current perceptual input and existing narrative models, triggering synchronized transitions across scales to update both perception and narrative understanding.

### Putative functional roles of scale-invariant anti-correlated activity patterns

Our findings reveal an interesting parallel between global and local anti-correlated activity patterns during naturalistic viewing. This parallel may not be coincidental but could reflect a scale-invariant organizational principle that serves similar cognitive functions across different spatial scales of neural organization. We propose that anti-correlated neural dynamics represent a fundamental mechanism for simultaneously maintaining continuity and enabling updating of information processing—at the global level supporting coherent narrative experience, and at the local level preserving specific contextual features within specialized processing regions.

#### Avoiding stimulus interference

Within the literature which specifically investigates anti-correlated patterns in local neural populations, a number of studies focus on how the brain segregates current and future goals within the same neural circuitry to prevent interference (***Collin et al., 2025***; ***Wan et al., 2020, 2022, 2024***). In ***van Loon et al. (2018***), the authors report that during tasks requiring memory of both current and prospective goals, the brain uses anti-correlated representational patterns in object-selective cortex regions to separate the two. Specifically, prospective/future goals are encoded in patterns that appear as an inverse of those for current/immediate goals. They suggest that this anti-correlation allows the brain to maintain prospective memory in a way that minimally disrupts the processing of ongoing tasks. These findings have since been supported by a number of other studies that have looked at encoding of currently and prospectively relevant items in working memory (***Collin et al., 2025***; ***Wan et al., 2020, 2022, 2024***; ***Yu et al., 2020b***). Generalizing their task-specific and non-continuous results to naturalistic settings, our findings also show that local populations switch between anti-correlated patterns when a novel ‘currently relevant’ stimulus feature is present. Thus we contribute additional credence to the theory that indeed stimulus interference might be avoided via anti-correlational patterns within which neural activity resides.

#### Maintaining Coherent Narratives and Local Context

At the global and local scale, the temporal dynamics and response to perceptual changes show similar patterns, suggesting a common underlying computational principle.

At the global level, anti-correlational switches may serve to maintain narrative coherence while enabling selective updating in response to meaningful changes in the environment. During periods dominated by the DMS, which engages regions associated with internal mentation, autobiographical memory, and semantic processing, the brain may be actively maintaining and elaborating a coherent narrative representation of ongoing experience. This narrative framework provides continuity and meaning to the stream of perceptual events. When salient perceptual changes occur—such as shifts in visual scenes, auditory features—the brain temporarily transitions to the TPS, focusing resources on processing these new inputs. Once this updating is complete, the return to DMS dominance may enable the integration of new information into the ongoing narrative structure, maintaining coherence across time despite changes in perceptual input. This oscillatory mechanism would allow the brain to strike a balance between stability and flexibility in narrative processing, maintaining a continuous thread of meaning while remaining responsive to new information.

At the local level, anti-correlated activity between adjacent voxel populations may implement a similar principle of simultaneous maintenance and updating, but specialized for the particular features processed in that region. While one population of voxels engages in processing immediate stimulus features (analogous to the global TPS), the anti-correlated population may maintain contextual information or representational templates (analogous to the global DMS) that situate current processing within broader temporal or semantic contexts. For instance, in visual regions, while one voxel population responds to current visual input, the anti-correlated population might maintain information about the visual context or recently processed visual features, enabling integration across time. Similarly, in language regions, anti-correlated populations could divide the labour between processing incoming linguistic input and maintaining the discourse context necessary for comprehension. This local anti-correlation would provide a mechanism for addressing what is known as the “stability-plasticity dilemma” in neural computation (***Grossberg, 1987***), i.e. how neural systems can remain sensitive to new information while preserving existing knowledge. By segregating these functions across anti-correlated neural populations, the brain could achieve both aims simultaneously within the same region, enabling efficient processing of new inputs while maintaining contextual continuity.

#### States of Mind

On the cognitive/experiential side, ***Herz et al. (2020***) propose a “States of Mind” framework, suggesting that mental states fluctuate along a continuum from broad to narrow state of mind, impacting perception, cognition, and affect. In this model, the DMN would align with top-down processes that support a broad state of mind, where perception and thought are expansive, integrative, and flexible. This broad state of mind is marked by exploratory thinking, openness to new ideas, and global attention—qualities that align closely with ***Yamashita et al. (2021***)’s optimal state for sustained focus, which also involves DMN activation and facilitates performance during a sustained attention task. Conversely, the narrow state of mind is driven more by bottom-up sensory processing and immediate task demands, leading to local attention and exploitative behaviour. This narrow focus corresponds with ***Yamashita et al. (2021***)’s suboptimal state, where there is low DMN activity and high TPN activity and performance tends to become less stable, with higher susceptibility to attentional lapses and distraction. ***Herz et al. (2020***)’s framework thus offers a cognitive perspective on how the DMN supports attentional states, not as a strictly “task-negative” network but as part of a system that shifts between broad and narrow states of mind depending on task complexity and motivational factors.

#### Computational principles and parallels with artificial intelligence

The anti-correlated neural states we observe across spatial scales bear resemblances to computational mechanisms which are useful for modern artificial neural networks (ANNs). Here we discuss a few computational mechanisms and how they might point to functional roles for anti-correlated states.

First, Recurrent Neural Networks (RNNs), and particularly gated architectures like Long Short-Term Memory (LSTM) networks (***Hochreiter and Schmidhuber, 1997***) and Gated Recurrent Units (GRUs) (***Cho et al., 2014***), are useful and performant models capable of time-series modelling. A key feature of these architectures is their ability to make use of gating mechanisms to selectively process new inputs while also (selectively) maintaining past information. This allows for complex processing which depends not only on recent but also on historical input data. This gating architecture has functional parallels to the anti-correlated neural dynamics we observe. The alternation between global DMS and TPS states resembles the operation of these gating mechanisms: TPS-dominant periods might be analogous to “input gate” activation, allowing new perceptual information to update representations, while DMS-dominant periods might function like “cell state” maintenance, preserving narrative continuity across time. A similar argument could be made for the separation of the internal model from the selective input-processing which exists within modern State Space models (SSMs) (***Gu and Dao, 2023***).

Taking another compatible example, the anti-correlated activity patterns which we observe, particularly at the local level where adjacent voxel populations show opposite response patterns, also evoke principles from contrastive learning in machine learning. Contrastive learning methods like SimCLR (***Chen et al., 2020***) and CLIP (***Radford et al., 2021***) learn representations by maximizing agreement between differently augmented views of the same data point (positive pairs) while minimizing agreement between representations of different data points (negative pairs). ***Konkle and Alvarez (2022***) demonstrated that self-supervised contrastive learning in artificial neural networks can lead to emergent organization of visual representations that resembles human brain responses, suggesting deep connections between contrastive learning principles and neural organization. The anti-correlated local patterns we observe could act as a substrate for such contrastive learning by maintaining opposing response patterns in adjacent neural populations such that they can not only be distinguished, but used for a continuous contrastive learning regime that enhances the discriminability of stimuli and their neural representations. This perspective aligns with ***van Loon et al. (2018***)’s finding that current and future goals are represented in opposite patterns in object-selective cortex.

Finally, the stimulus aligned transitions which we observe also connect to predictive coding theories of brain function (***Rao and Ballard, 1999***; ***Friston, 2010***), which propose that the brain constantly generates predictions about incoming sensory information and computes prediction errors when these expectations are violated. In predictive coding frameworks, prediction errors drive updates to internal models, allowing the brain to refine its predictions over time. Our finding that transitions from DMS to TPS are often triggered by salient perceptual changes could indicate that these are moments of heightened prediction error processing, during which the brain updates its internal models based on unexpected or sudden novel sensory information. The subsequent return to DMS may reflect the incorporation of these updates into refined predictive models that can generate more accurate predictions moving forward. Such a mechanism could occur across scales with local anti-correlated voxel populations implementing a similar predictive coding mechanism at a more local scale.

This set of computational principles are an interesting set of initial hypotheses with which one could explore the underlying functional role of the anti-correlated neural state patterns and their relation to ongoing stimulus processing.

### Limitations and Future research directions

First, we find it important to discuss and clarify one of the main terms used throughout the paper: anti-correlated. Although we observe anti-correlated activity patterns in our data, a number of different neural activity structures could have given rise to these observations. These alternative structures of neural dynamics would have implications for exactly how and what is being encoded within underlying neural populations. To be specific, the signals observed herein could be the result of a continuous pattern, where neurons are spiking along a continuum or range of intensities, but with an anti-correlated pattern between two populations. Alternatively, these signals could reflect a binarised pattern, where there is a strong competitive or antagonistic dynamic such that only one subdivision of the population tends to be active at a time. The latter condition would appear as if these two populations separately encode information but tend not to be coactive. This is equivalent to the two populations encoding in orthogonal spaces, but specifically where only one population (space) is typically occupied at any moment. Given such a possibility, we caution any readers from over-interpretting the label of ‘anti-correlation’, as this could lead to mis-characterisations of the underlying phenomena driving these dynamics. We leave the investigation of the consequences of these different interpretations to future work.

Future research should also investigate the temporal order in which state transitions happen between local ROIs and/or the global brain. Here, we only present overlapping transitions, but studying states which consistently transition before or after one another could shed further light into the information flow across the cortex in naturalistic settings. Neural states and patterns which we labelled as the univariate states also warrant further investigation. As more idiosyncratic periods of “reconfiguration”, they likely are behaviourally relevant. Last but not least, the role of these anti-correlated/antagonistic/orthogonal states (across scales) in representing information should also be further studied. Understanding how neural representations evolve over time through anti-correlated states could inform us about mechanisms behind cognitive processes such as working memory, attention, and abstraction.

### Conclusion

In this study, we have demonstrated that brain activity during naturalistic viewing is dominated by two anti-correlated states that oscillate in a semi-regular fashion, with a third transitional state mediating between them. This organization exists at both global and local spatial scales, with DAN regions such as the frontal eye fields and superior parietal lobule showing particularly strong temporal alignment between scales. We found that transitions between states are systematically triggered by changes in perceptual input in a modality-specific manner, with visual changes driving transitions in visual cortical regions and auditory changes affecting auditory and language-related areas. These findings were robust across multiple datasets with different stimulus types, indicating that anti-correlated neural dynamics represent a domain-general organizational principle of brain function.

The anti-correlated states we observe likely implement a fundamental operating rhythm for naturalistic cognition, alternating between periods of externally-focused information processing (TPS) and internally-directed integration and maintenance (DMS). This rhythmic alternation may provide a neural mechanism for simultaneously maintaining narrative continuity while remaining responsive to meaningful changes in the environment, which is a core requirement for adaptive cognition in natural settings. The scale-invariant nature of these dynamics suggests they may represent a core computational motif that enables flexible yet stable information processing across different levels of neural organization. Global anti-correlation has even been suggested to play a central role in consciousness (***Demertzi et al., 2022***).

By revealing the spatiotemporal principles that govern naturalistic brain activity, our findings advance our understanding of how the brain constructs coherent experience from the continuous stream of sensory input we encounter in everyday life. Ultimately, the anti-correlated neural states we have identified may represent a fundamental organizing principle of brain function that underlies our ability to construct coherent, continuous experience from the stream of perceptual events we encounter in the world. This ability to maintain narrative continuity while selectively updating our understanding in response to meaningful changes lies at the heart of human cognition and consciousness.

## Contributions

**Dora Gözükara**: Conceptualization, Methodology, Software, Formal analysis, Visualization, Writing - Original Draft. **Djamari Oetringer**: Methodology, Data Pre-processing, Software, Writing - Review. **Nasir Ahmad**: Methodology, Software, Formal analysis, Visualization, Writing - Original Draft & Review & Editing, Supervision. **Linda Geerligs**: Conceptualization, Methodology, Validation, Writing - Original Draft & Review & Editing, Funding acquisition, Supervision.

## Funding

Linda Geerligs was supported by a Vidi grant (VI.Vidi.201.150) from the Netherlands Organization for Scientific Research.

## Declaration of Competing Interests

The authors declare that no competing interests exist.

## Acknowledgments

We would like to thank Emily Mo Nipshagen for useful deliberations on the early statistical analyses; Mesian Tilmatine for insights in narrative comprehension; and Sahel Azizpour for constructive discussions at every stage of the paper. We thank by Karen Campbell and Chloe Winters for providing the annotations for the CamCAN movie.

## Methods and Materials

### Datasets

#### Study Forrest

We used the open access StudyForrest fMRI dataset in which 15 participants watched the movie Forrest Gump dubbed in German (age 19–30, mean 22.4, 10 female, 5 male) (***Hanke et al., 2016***). The 2-hour movie was divided into 8 segments that were presented across two sessions on the same day. In total there were 3599 volumes of data acquired with a 32-channel head coil using a whole-body 3 Tesla Philips Achieva dStream MRI scanner. The acquired images were T2*-weighted echo-planar images (gradient-echo, 2 s repetition time (TR), 30 ms echo time, 90° flip angle, 1943 Hz/Px bandwidth, parallel acquisition with sensitivity encoding (SENSE) reduction factor 2). There were 35 axial slices (thickness 3.0 mm) with 80 × 80 voxels (3.0 × 3.0 mm) of in-plane resolution, 240 mm field-of-view (FoV), and an anterior-to-posterior phase encoding direction with a 10% inter-slice gap, recorded in ascending order. A T1-weighted image was acquired for anatomical alignment (***Hanke et al., 2014***).

#### Preprocessing

We used data that was partially preprocessed by ***Liu et al. (2022***). They corrected the data for motion and slice timing, extracted the brain, and applied a high-pass temporal filter of 1 Hz. They also denoised the data by using spatial ICA decomposition and manual inspection. We used SPM12 (***Wellcome Centre for Human Neuroimaging, 2014***) to perform a number of additional pre-processing steps. These pre-processing steps involved: aligning the functional data between each run, co-registering the data to the anatomical volumes of each participant, and normalising the data to MNI space. Afterwards, we z-scored the data per voxel per run.

In our analyses, we only included grey matter voxels that were active in all runs. Specifically, the gray matter mask contained voxels that survive the threshold of 0.35 in the gray matter probability map (TPM) as provided by the SPM software. We defined voxels as active per run if their response at the first timepoint was higher than 0.8 times the mean of all voxels at that same timepoint. To optimize the alignment of data across participants, we used whole-brain searchlight hyperalignment, using the PyMVPA toolbox (***Haxby et al., 2011***) implementation. We used the fourth run (488 volumes) to compute the hyperalignment parameters and applied the parameters to all remaining runs. The fourth run was not included in any subsequent analyses.

#### Stimulus Annotations

We used the shot transitions in the movie to determine changes in visual features as defined by ***Häusler and Hanke (2016***). To determine auditory features we used moment to moment changes in Mel-Frequency Cepstral Coefficients (MFCC) of the audio, as computed by ***Oetringer et al. (2025***). Event boundaries were defined by ***Ben-Yakov and Henson (2018***), who had independent raters determine changes in abstract narrative events. All annotation types except the MFCC change were binary indicators of a shot/event changes. MFCC annotations were cosine distances between MFCC patterns in consecutive timepoints (downsampled to 2 seconds to match the TR), which were later binarised by setting all timepoints which were greater than one standard deviation from the mean to one, and the rest to zero. All annotations were delayed by 4.5 seconds to correct for the haemodynamic response delay, as computed by ***Oetringer et al. (2025***) in an independent analysis.

#### CamCAN

We used the data from 265 adults (131 females) who were aged 18–50 (mean age 36.3, SD = 8.6) from the healthy cohort recorded in the stage II of the Cam-CAN project (***Shafto et al., 2014***; ***Taylor et al., 2017***). Participants were native English speakers, had normal or corrected-to-normal vision and hearing, and had no neurological disorders. Participants viewed an edited version of Alfred Hitchcock’s black-and-white television narrative “Bang! You’re Dead” (***Hasson et al., 2008, 2010***) edited from 30 min to 8 min while maintaining the plot (***Shafto et al., 2014***). In total there were 192 volumes of data acquired with a 32-channel head coil using a 3T Siemens TIM Trio scanner at the Medical Research Council Cognition and Brain Sciences Unit. The acquired images were multi-echo, T2*-weighted echo-planar images (TR = 2470 ms, five echoes [TE = 9.4, 21.2, 33, 45, 57 ms], flip angle = 78°, v3 × 3 × 4.44 mm voxel size). There were 32 axial slices (thickness 3.7 mm) with a 20% inter-slice gap, 192 × 192 mm field-of-view (FoV), recorded in interleaved order. A high-resolution T1-weighted image was acquired for anatomical alignment using a magnetization prepared rapid acquisition gradient-echo (MP-RAGE) pulse sequence (1 × 1 × 1 mm resolution, TR = 2250 ms, TE = 2.99 ms, TI = 900 ms, flip angle = 9°, FOV = 256 × 240 × 192 mm, GRAPPA acceleration factor = 2).

#### Preprocessing

The preprocessing of the data was identical to previous work with the same dataset (***Geerligs et al., 2022***), we will briefly reiterate the main steps here. Preprocessing included deobliquing of each TE, slice time correction, and realignment of each TE to the first TE in the run, using AFNI (***Cox, 1996***). Multi-echo independent component analysis (ME-ICA) was used to denoise the data (***Kundu et al., 2012***). Functional data were co-registered to the anatomical scans and DARTEL intersubject alignment was used to align participants to MNI space using SPM12 software (***Wellcome Centre for Human Neuroimaging, 2014***). To optimize the alignment of voxels across particiapnts, whole-brain searchlight hyperalignment was used, as implemented in the PyMVPA toolbox (***Haxby et al., 2011***). After hyperalignment, the data were highpass-filtered with a cut-off of 8 millihertz (mHz). Afterwards, we z-scored the data per voxel.

#### Stimuli Annotations

We used event boundaries defined by independent raters to determine changes in abstract narrative events acquired by ***Ben-Yakov and Henson (2018***). In addition, we used annotations coded according to the guidelines of ***Zacks (2010***). The annotations comprised nine categories of event boundaries: Character boundaries occurred when the focus of attention shifted to a different animate character; CharChar boundaries marked changes in physical or abstract interactions between characters (e.g., touching, talking, gesturing); CharObj boundaries indicated changes in a character’s interaction with objects (e.g., picking up, putting down, or using an object differently); Time boundaries denoted temporal discontinuities following cuts; SpaceLg boundaries marked transitions to new spaces or locations; SpaceSm boundaries occurred when characters changed direction of motion within a scene or when camera viewpoint changed location; Cause boundaries indicated when activity in one frame could not be directly explained by activity in the previous frame; Goal boundaries marked when characters performed actions associated with different goals compared to the previous frame; and Scene boundaries indicated transitions to new scenes. Due to the way these are defined, there was a lot of overlap between boundaries defined across these categories. All boundary annotations were binary indicators of event changes and were delayed by a 4.5 seconds haemodynamic response delay.

#### Narrattention

##### Story Listening

54 participants (mean age of 23, 40 female, 14 male) listened to 3 Dutch stories taken from the audiobook series “Joyride & andere spannende verhalen” by René Appel. In particular, the selected stories were “Vader en Zoon” (*Father and Son*), “Boekenliefde” (*Book Love*), and “Heterdaad” (*Red-handed*). The original audiobooks ranged from 11 to 15 minutes, but were cut down to approximately 10 minutes each during stimulus presentation. A fixation cross was presented on the screen, but without any explicit instructions on actually fixating on the cross, to prevent participants from having an additional non-naturalistic task. Participants were also allowed to close their eyes during the stories, as long as this did not make them tired. This dataset was collected in the context of another study, in which an attention manipulation was applied. As part of this attention manipulation, the auditory stories were edited such that there were two narrators taking turns, and the stimulus would be presented dominantly in the left ear at some moments and dominantly in the right ear at others. Each participant was presented with the exact same stimulus with identical edits, but were given different instructions for the same stimulus: some were told to attend the narrator by keeping track of who spoke more, others were instructed to attend the sound location by keeping track of in which ear the stimulus was presented the longest, and again others were told to just listen to the stories. Here we only used those parts of the data where participants were instructed to simply listen to the stories, meaning that they did not have an extra attention task on top of listening to the story. As the design of the attention conditions were within-participant, we have a different set of participants for each story (18 participants per story).

##### Movie-watching

The same set of participants who listened to the stories also viewed an audiovisual movie of roughly 10 minutes, which was a cut-down version of a 45-minute episode of a Dutch detective television series. They were instructed to watch the movie as they would at home, and thus they were free to move their eyes around. This data was used to hyperalign the story listening data. In this study, the non-hyperaligned movie data was also used to create the local template patterns map to verify that our results were not a product of hyperalignment alone (see 2).

##### Preprocessing

All BOLD runs used in this story (3 story listening runs and 1 movie watching run described below) and the T1-weighted anatomical image were preprocessed using *fMRIPrep* 23.0.2 (***Esteban et al., 2019, 2018***), which is based on *Nipype* 1.8.6 (***Gorgolewski et al., 2011, 2018***). The T1-weighted (T1w) image was corrected for intensity non-uniformity (INU) with N4BiasFieldCorrection (***Tustison et al., 2010***), distributed with *ANTs* 2.3.3 (***Avants et al., 2008***), and used as T1w-reference throughout the workflow. Volume-based spatial normalization to one standard space (*MNI152NLin2009cAsym*) was performed through nonlinear registration with antsRegistration, using brain-extracted versions of both T1w reference and the T1w template.

For the BOLD runs, first a reference volume and its skull-stripped version were generated by aligning and averaging 1 single-band references (SBRefs). Head-motion parameters with respect to the BOLD reference (transformation matrices, and six corresponding rotation and translation parameters) are estimated before any spatiotemporal filtering using *mcflirt* (*FSL* 6.0.5.1, ***Jenkinson et al. (2002***)). BOLD runs were slice-time corrected to 0.696s (0.5 of slice acquisition range 0s-1.39s) using *3dTshift* from *AFNI* (***Cox and Hyde, 1997***). The BOLD time-series were resampled onto their original, native space by applying the transforms to correct for head-motion. The BOLD reference was then co-registered to the T1w reference using *bbregister* (FreeSurfer) which implements boundary-based registration (***Greve and Fischl, 2009***). Co-registration was configured with six degrees of freedom. First, a reference volume and its skull-stripped version were generated using a custom methodology of fMRIPrep. The BOLD time-series were resampled into standard space, generating preprocessed BOLD runs in *MNI152NLin2009cAsym* space. Additionally, ICA components were computed and labelled as motion-noise or not-motion-noise using ICA-AROMA (***Pruim et al., 2015***).

The remaining preprocessing pipeline was implemented in *Python*. First, motion artefacts were removed from all BOLD runs by applying the ICA components and ICA-AROMA labels using a non-aggressive correction (***Pruim et al., 2015***). White matter and cerebrospinal fluid signal were removed non-aggressively as well, followed by gray matter masking. Here, the gray matter was defined as voxels that were measured for each participant and had a value of at least 0.35 in in the gray matter probability map (TPM) as provided by the SPM software. Then, all BOLD runs were high-pass filtered using a discrete cosine transform with a cut-off of 200 seconds, and the data was z-scored to give a mean activity of zero per voxel.

Finally, the data was hyperaligned by computing the parameters using the movie run, and applying these parameters to all remaining runs. Hyperalignment functionally aligns the participants’ data to each other, improving inter-subject correlation. Specifically, we used searchlight-hyperalignment using the implementation of the PyMVPA toolbox (***Guntupalli et al., 2016***; ***Haxby et al., 2020***) in which each searchlight had a radius of 3 voxels and all voxels within the gray matter mask were used as the center of a searchlight. Then some starting and ending volumes were removed as they contained buffers and spoken recall questions. Finally, the remaining data was z-scored again, as hyperalignment and the cutting of the data moves timeseries means away from zero.

##### Stimuli Annotations

This dataset included annotations for narrative event boundaries. To extract event boundaries from the stories, an experiment with an independent set of participants (N = 33, 21 female, 12 male, aged 19 to 59 years old, with an average age of 25.8 years) was performed. After explaining the concept of event segmentation, participants were instructed to press a button whenever they thought a new unit had started. Half of the participants were instructed to make the units as big as possible while still being meaningful, measuring coarse-grained event boundaries. The other half was instructed to make the units as small as possible while still being meaningful, measuring fine-grained event boundaries. In this study, we only used the coarse-grained results. Button presses were taken together between participants by first shifting all presses to the start of a phrase, and then defining those moments as a boundary if the number of participants pressing the button exceeded a certain threshold. Similar to ***Ben-Yakov and Henson (2018***), the threshold was determined per story per condition by first averaging the total number of button presses across participants, and setting the threshold such that the number of boundaries was closest to these averages, giving a threshold of 4 or 5 participants depending on the story.

### Data Processing

#### Schaefer Atlas

For all datasets, we made use of the Schaefer Atlas ***Schaefer et al. (2017***) with 400 distinct regions of interest (ROIs) distributed across 17 functional networks. This parcellation is based on resting-state functional connectivity data and is provided in the Python nilearn package (***Abraham et al., 2014***). We obtained the Schaefer 400-ROI, 17-network atlas from nilearn and resampled the atlas from its native MNI152 space to match the dimensions and voxel size of our fMRI volumes (also in MNI152 space). We extracted data from each voxel within each of the 400 ROIs in the atlas, excluding any voxels that were not part of the gray matter or not active in each run.

#### Brain Data

Each ROI, indexed *k* ∈ [1, 2, …, *K*] with *K* = 400, represents a distinct brain region in the Schaefer atlas with a corresponding network assignment.

Let us define the following notations:

- *K* = 400: The total number of ROIs in the Schaefer atlas
- *T* : The number of time points in the fMRI acquisition
- *V*_*k*_: The number of voxels in ROI *k*
- 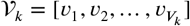: The set of voxels belonging to ROI *k*, of length *V*_*k*_

Using these notations, we define:

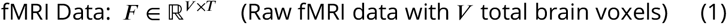

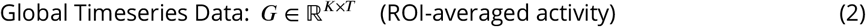

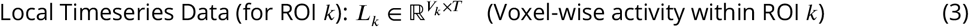

For each *k*th ROI from the Schaefer atlas and time point *t*, the mathematical relationship between the local and global timeseries is:

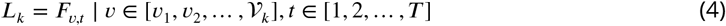

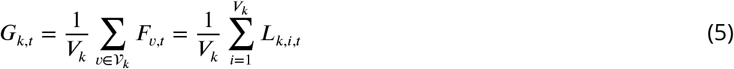

This formulation shows that the “Global Timeseries” is derived by averaging voxel-wise activity within each Schaefer atlas ROI, and that the *k*th “Local Timeseries”represents the original fMRI time series data for all voxels within ROI *k*.

#### Correlation Matrices

To identify and characterize anti-correlated states in multivariate fMRI data, we first computed the temporal correlation matrix of the multivariate timeseries (either voxel by time or ROI by time) to capture similarities between timepoints. We then calculated a meta-correlation matrix to reveal higher-order relationships between correlation patterns (for an interesting new approach, see ***Hindriks et al. (2024***)).

Specifically, we can describe our local ROI or global timeseries data as a 2D array of multivariate timeseries data *X* ∈ ℝ^*N*×*T*^, where *n* is the number of features and *t* the number of timepoints. Given this multivariate timeseries data *X*, the time-by-time correlation matrix *C* ∈ ℝ^*T* ×*T*^ can be expressed as:

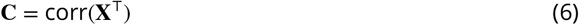

where each individual element of this matrix represents the Pearson correlation coefficient between the spatial patterns at timepoints i and j:

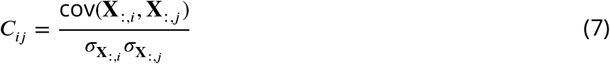

where cov(⋅) indicates the covariance between two variables, and σ_⋅_ indicates the standard deviation of a variable.

With this definition, we then calculate a meta-correlation matrix to reveal relationships between the correlation patterns *within* the correlation matrix:

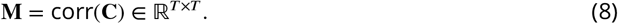

Finally, we then applied Hartigan’s dip test to each row of the meta-correlation matrix **M**, getting both the dip statistic (measuring deviation from unimodality) and its associated p-value (where lower values indicate stronger evidence for multimodality) (***Urlus, 2023***; ***Hartigan and Hartigan, 1985***; ***Hartigan, 1985***).

#### Test of Stationarity

To assess the stationarity of the timepoint correlation structure, we performed Augmented Dickey-Fuller (ADF) tests on each timepoints correlation pattern across the timeseries. The ADF test was applied to both whole-brain and region-of-interest (ROI) correlation data, using the adfuller() function from Python’s statsmodels package (***Seabold and Perktold, 2010***). For each timepoint, we extracted its correlation vector with all other timepoints and tested the null hypothesis of a unit root (non-stationarity) against the alternative hypothesis of stationarity (***MacKinnon, 1994***). P-values from the ADF tests were collected for all timepoints in both the whole-brain and ROI datasets.

#### State and Template Definitions

Neural state templates were created from multivariate data using only timepoints which exhibit bimodal meta correlation distributions. To identify these bimodal datapoints we applied Hartigan’s dip test to every row of the meta-correlation matrix to arrive at a set of p-values, one for each time-point *p*^dip^ ∈ ℝ^*T*)^. Using these p-values, timepoints exhibiting bimodal distributions 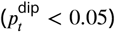 were extracted, 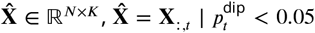 where *K* is the number of selected (bimodal correlation structure) timepoints. To be able to identify a pattern that is either negatively or positively expressed at different time points, we first adjust the sign of all timepoints. such that anti-correlated timepoints are reversed. In practice, we correlate each timepoints with the first timepoints and then sign flip any timepoint that is negatively correlated with that first timepoint. This pipeline defined our template patterns.

In all, this can be expressed such that,

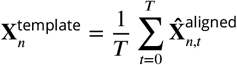

where

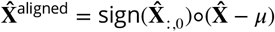

where ° indicates Hadamard product (element-wise multiplication of the vector elements with every element in the matrix) and 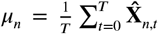 describes the elements of the demeaning vector. Note that this process is equivalent to a linear regression over our bimodal timepoints in which they are each assigned either a ‘+1’ or ‘-1’ weighting depending upon whether they are sign aligned or mis-aligned with the first timepoint’s correlation structure.

This way, we captured the dominant pattern of coordinated activity across features which corresponded to the major axis of while accounting for anti-correlated flips.

Finally, we mapped the multivariate data *X* ∈ ℝ^*N*×*T*^ onto the template space to quantify the expression of the template pattern at each timepoint. To do so we first centred the data by calculating **X**^centered^ = **X** − ***µ***°**1**. The projection coefficients are then computed as:

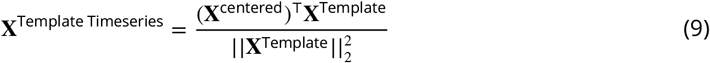

where **X**^Template Timeseries^ ∈ ℝ^1×*T*^ represents the strength and direction of template expression at each timepoint, with positive values indicating alignment with the template pattern and negative values indicating alignment with the opposite pattern. To make sure that the sign of the template was meaningful, we aligned it to the mean activity in the data. If the mean activity was negatively correlated with the template timeseries, we flipped the sign of the template and recomputed the timeseries. This can be expressed as

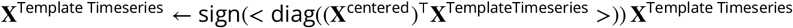

where sign(⋅) is the sign function, and diag(⋅) returns the diagonal entries of a matrix.

#### Proportion of Variance Explained by the Templates

To assess the quality of the global template reconstruction, we calculated the proportion of variance explained as 1 − (MSE/σ^2^), where MSE represents the mean squared error between the original ROI-averaged data and the global template timeseries, and σ^2^ represents the total variance in the original data. Both signals were mean-centred prior to calculation. This metric quantifies how much of the original signal’s variance structure is preserved in the template, with values of 1.0 indicating perfect reconstruction and 0.0 indicating reconstruction quality equivalent to the mean.

#### Mean BOLD Activity Difference Between Local Anti-correlated States

To test for significant mean BOLD activity differences local anti-correlated states, we performed Welch’s t-tests (assuming unequal variances) comparing the mean activities of timepoints that are labelled as the two different states across all ROIs. To correct for multiple testing, we applied false discovery rate control (FDR) across across ROIs.

### Local-Global State Transition Alignment

Global state transitions were identified as points where the sign of the global template projection changed, and similarly local state transitions were identified as points where the sign of the local template projection changed. We defined a match by global and local transitions when transitions occurred at the same timepoint. In order to quantify whether the overlap between neural state boundaries across local and global scales was larger than expected, we computed the absolute boundary overlap metric as described in ***Geerligs et al. (2022***). The absolute boundary overlap metric (OA) was scaled such that it was zero when it was equal to the expected overlap and one when it was equal to the maximum possible overlap. The absolute overlap metric (OA) measures overlap between two different ‘boundary’ annotations by using the minimum number of boundaries of either type as its reference, and normalizing the difference between this minimum and the expected overlap.

### Relation to Stimuli

To quantify the relationship between global neural states and stimulus boundaries, we used a permutation-based statistical test. At each timepoint we know whether the brain was in the TPS or DMN state (at the global scale) or the up or down state (at the local scale). To compute the probability of a timepoint around a stimulus boundary being in a specific state, we first express this specific state as a 1 and the other state as 0, before averaging the values across all stimulus boundaries. This gives us a metric of the state probability at each timepoint around stimulus onset. Next, we compared this to a null distribution generated through permutation testing. The null distribution was created by preserving the durations of states while randomly shuffling their order across 100,000 iterations for the global scale and 10,000 for the local scale. This analysis was applied systematically across all stimuli types (events, shots, MFCC and other annotations) and neural states. For the global scale, this was done within a temporal window ranging from -10 to +10 timepoints to capture effects in time. For the local scale, we used a temporal window ranging from -3 to +3 timepoints. As a result, for each stimulus and state transition type, we calculated p-values and differences between how often states occur on average and how often they occur around stimuli boundaries. To correct for multiple testing, we applied false discovery rate control (FDR) across timepoints (for the global scale) and across ROIs (for the local scale). Note, due to our definition of only two main states (globally DMS/TPS and locally up/down), any change in transition probability from one state to the other has a corresponding (and exact) opposite impact upon transition probability in the opposite state transition direction. Therefore all measures of state change probability are mirrored both visually and statistically.

### Combining results across Narrattention stories

Unlike the other datasets, where all participants were subject to the same stimuli, Narrattention had three stories which were listened by different groups of participants. We therefore adjusted our statistical analysis to be able to combine effects across these three stories.

For calculating the global and local spatial template patterns, as well as the overlap between global and local state boundaries, we simply averaged the template patterns and boundary overlap maps across all stories.

To test the relationships between neural states and event boundaries, we adjusted the permutation-based statistical framework we used to pool evidence across all three stories within this dataset.

Specifically, we computed the probability of each of the global and local states at different time-points around event onset (like we described above) for each story separately. Then we averaged these probabilities across the three stories. To obtain null distributions, permutation tests were also performed as described above. We then pooled the permutation distributions and observed overlap values across all three stories for each temporal delay. This was done by averaging the probabilities across the three stories for each permutation. The resulting p-value was calculated as the proportion of pooled null values exceeding the average observed value. Subsequently false discovery rate correction for multiple comparisons was performed as described above.

### Cortical Mapping and Surface Projections

In all plotted cortical maps, except the visualisation of the local templates, we have projected the results of each ROI onto all the voxels they cover. For the local template visualisation, individual voxels within ROIs had unique values. We mapped the volumetric results on an average inflated cortical surface mesh (fsaverage) for our figures. All volumetric nifti files are also available with the code.

## Data and code availability

The entire analysis pipeline can be found at https://github.com/drgzkr/Multi-Scale_Anti-Correlated as a Jupyter Notebook. The data we used are also available at the group average whole-brain stage. The notebook explains how to use your own data (you will need a group-averaged, whole-brain (preferably hyperaligned, but not necessarily) nifti file), and stimuli annotations if you have any.

## Appendix 1

### Appx.1: DMS/TPS Annotation

Figure 1 shows an anatomical and functional overview of the global anti-correlated states across the whole brain.

**Appendix 1—figure 1.**
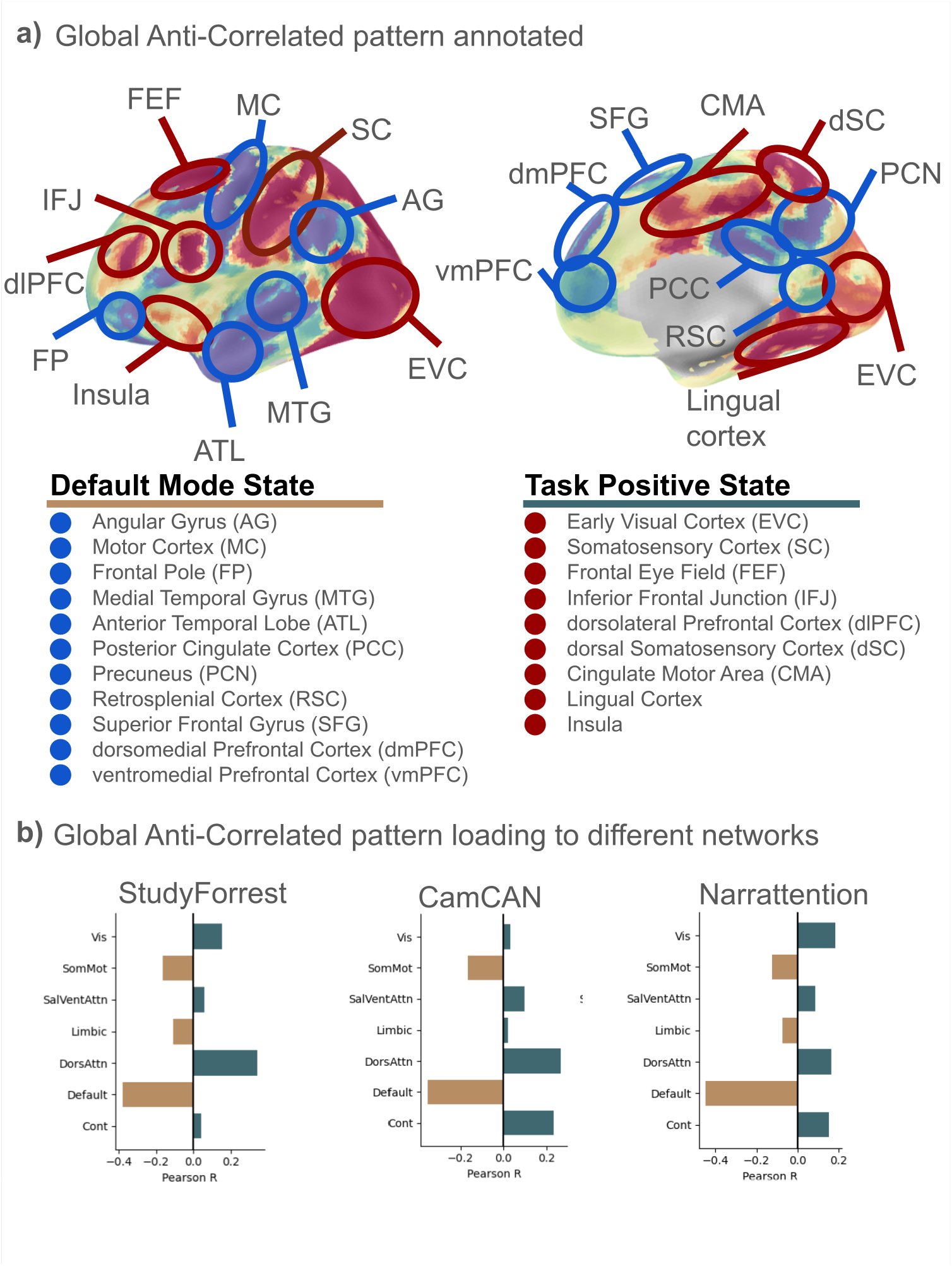
Key brain areas and networks annotated with the global anti-correlated templates. a) The annotated cortical map (same map in 2a) shows which ROIs participate in which of the global anti-correlated states. Blue shows the default mode state and red shows task positive state. b) Correlation between the each of the functional brain networks defined by ***Thomas Yeo et al. (2011***) and global anti-correlated state for each dataset. Brown is default mode state and blue is task positive state.

## Appendix 2

### Appx.2: Template patterns in more detail

To verify the spatial pattern of the anti-correlated templates in our results, we created a voxel-wise whole brain template (using the StudyForrest) and the local template patterns from non-hyperaligned data for two of our datasets. The voxel-wise whole brain template is created by using all the measured voxels across the cortex without any averaging or masking. As a result (and in contrast with the local template patterns show in other figures), we can see which global anti-correlated state a voxel is more associated with, but also the relative anti-correlated relationship between neighbouring voxels at different colour bar ranges. This map confirms that the local anti-correlation is still present even when we don’t reference voxel activity to a local context. We also see that the fine mesoscale organisation seen in the local template patterns with hyperaligned data is still present in non-hyperaligned data, but the structure is coarser with bigger anti-correlated patches. This is likely because just the anatomical alignment between participants averages detailed activity across participants leaving only coarser shared responses, while hyperalignment preserves finer patterning, albeit in a slightly transformed local anatomical space. This indicates that our findings are not a by-product of hyperalignment, but hyperalignment indeed enables us to distinguish finer spatial details.

**Appendix 2—figure 1.**
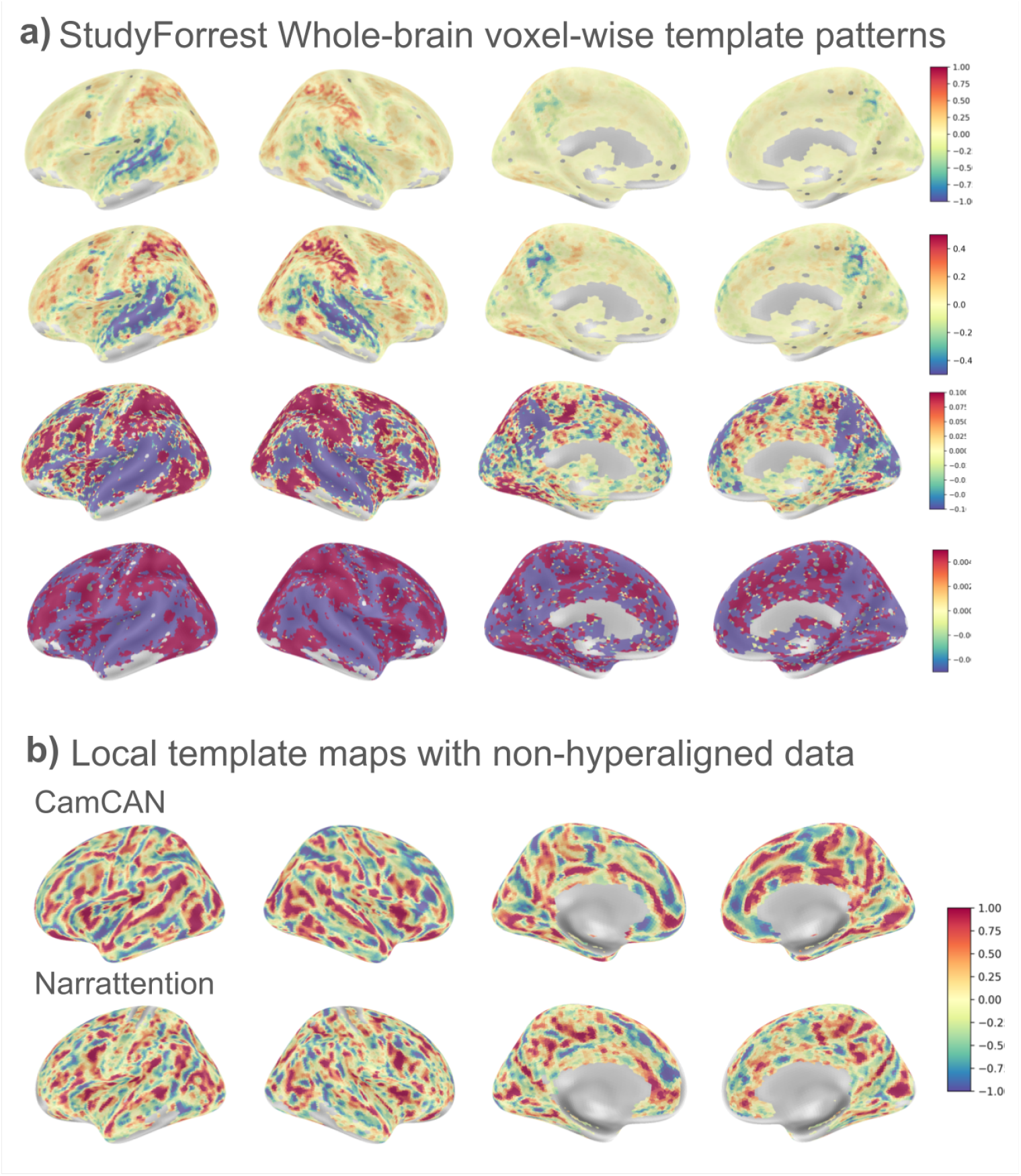
Local template patterns in more detail. (a) Voxel-wise Global Template. Each row shows the same data but with different colour map ranges to highlight the contrast within and between local and global scales. (b) Two example voxels from a visual ROI shows how neighbouring voxels show anti-correlated activities. (c) Local template patterns computed from non-hyperaligned data.

Figure2 shows some voxel timeseries for example ROIs and the time by time correlation matrix of the ROIs multivoxel timecourse.

**Appendix 2—figure 2.**
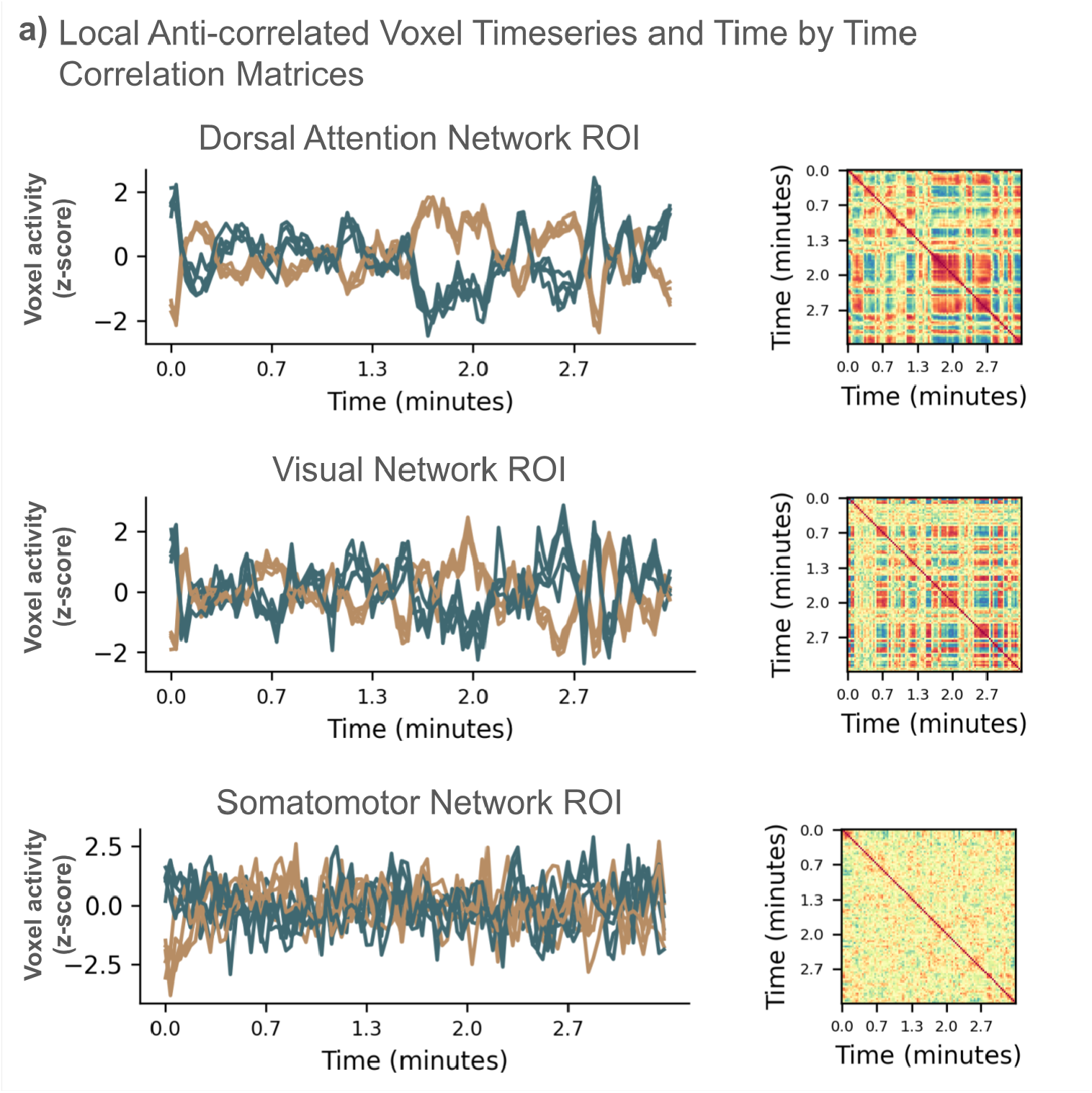
Anti-correlated voxel timeseries and time by time correlation matrices for example ROIs. Here, we show data from three example ROIs. For each, 10 voxels were selected from extreme ends of the template pattern to illustrate the local anti-correlation. These three specific ROIs were selected to illustrate the range of the degree of anti-correlation across ROIs. Although most ROIs contain clearly anti-correlated groups of voxels, the degree of anti-correlation is much less prominent for some ROIs.

## Appendix 3

### Appx.3: Relation to stimuli for CamCAN dataset

We performed the same stimuli relation analyses we presented for StudyForrest in the results with various stimuli annotations available for the CamCAN dataset. We have decided to only investigate the alignment with annotations which had less transitions than half the timepoints in the fMRI data (192 total volumes, thus annotations with fewer than 96 instances). This left us with event boundary annotations (independent rater coded, 19 boundaries), changes in time (frames after a cut is temporally discontinuous with the frame preceding the cut, 12 boundaries), cause (when activity in one frame cannot be directly explained (caused) by an activity viewed in the previous frame (e.g., character initiating a new action), 26 boundaries), and goal (where the character performs an action associated with a goal different to that of the previous frame (e.g., when a character initiates speaking), 52 boundaries).

Results show that only changes in goals are statistically significantly associated with a global brain state, and it is the default mode state that is more likely to coincide a goal switch, not the task positive state.

**Appendix 3—figure 1.**
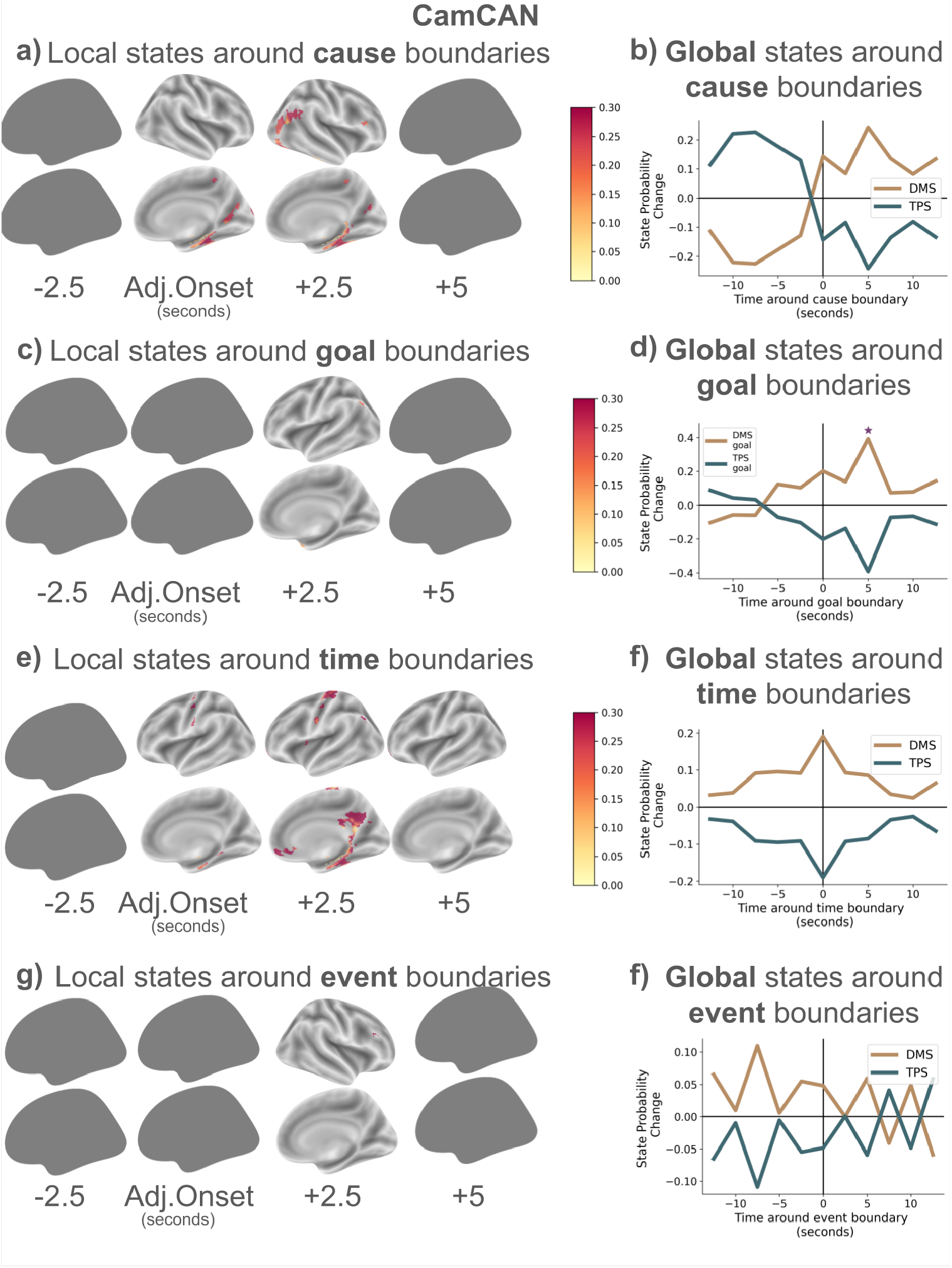
Relationship between various stimuli boundaries and local and global states for CamCAN. (a) Increase in probability of being in the up state spreads across the cortex after cause boundaries. (b) Probability of being in the TPS increases after cause boundaries. (c) Increase in probability of being in the up state compared to baseline spreads across the cortex after goal changes. (d) Probability of being in the TPS increases after goal boundaries. (e) Increase in probability of being in the up state spreads across the cortex after time changes. (f) Probability of being in the TPS increases after time boundaries. Results are visualized as state probability changes around boundaries, with significant timepoints (p<0.05 after FDR correction) marked with asterisks. Empty silhouettes show that no statistically significant ROIs are found.

## Appendix 4

### Appx.4: Relation to stimuli for Narrattention dataset

We performed the same stimuli relation analyses we presented for StudyForrest in the results with event boundary annotations available for the Narrattention dataset.

**Appendix 4—figure 1.**
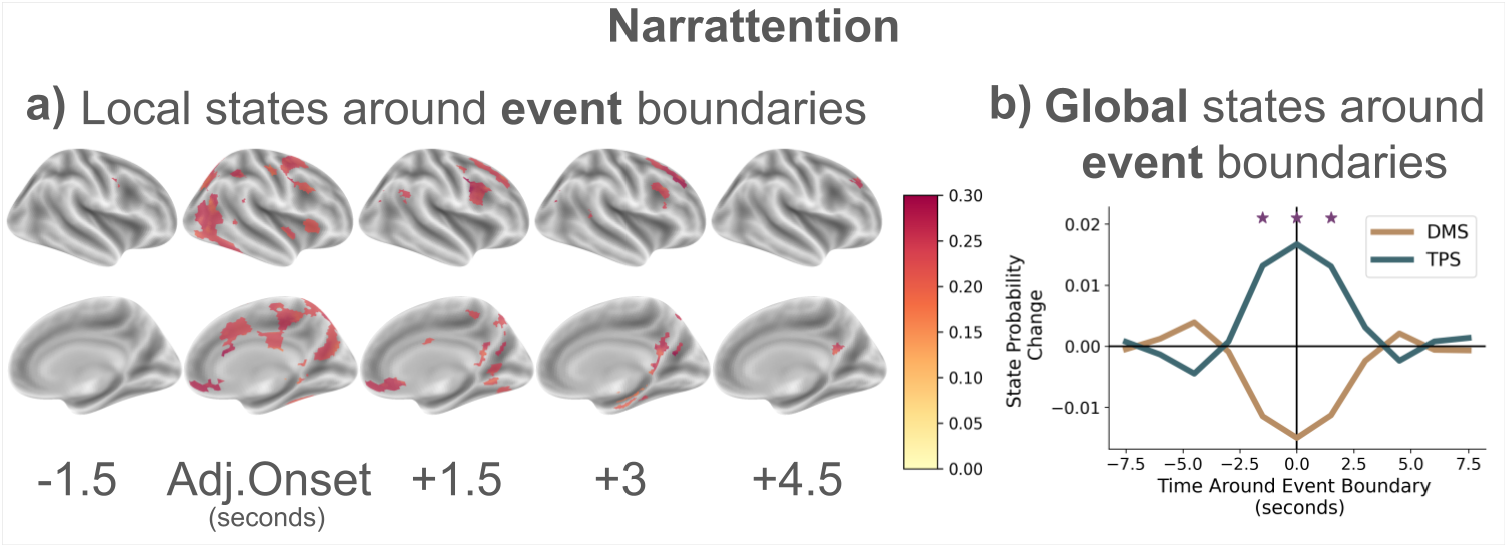
Relationship between various stimuli boundaries and local and global states for Narrattention. (a) Increase in probability of being in the up state spreads across the cortex after event boundaries. (b) Probability of being in the TPS increases after event boundaries. Results are visualized as state probability changes around boundaries, with significant timepoints (p<0.05 after FDR correction) marked with asterisks

## Appendix 5

### Appx.5: Global-Local state boundary overlap results across datasets

The sets of brain areas whose local state transitions align with the global state transitions show differences across datasets. In particular, we observe that We attribute these differences to the differences in the properties of the different stimuli which were used. The StudyForrest dataset has around two hours (3000+ volumes with a TR of 2 seconds) of fMRI recordings from 15 participants watching a minimally edited movie. The CamCAN dataset offers a much shorter duration with about eight minutes (192 volumes with a TR of 2.5 seconds) per participant, but has around 250 participants. The stimuli is black and white, and is a heavily edited (shortened) version of an already short (fifteen minutes) movie. The Narrattention dataset contains three audio narratives (short stories) each around ten minutes in length (407, 415, and 420 volumes with a TR of 1.5 seconds), with a constant gray background on the screen.

**Appendix 5—figure 1.**
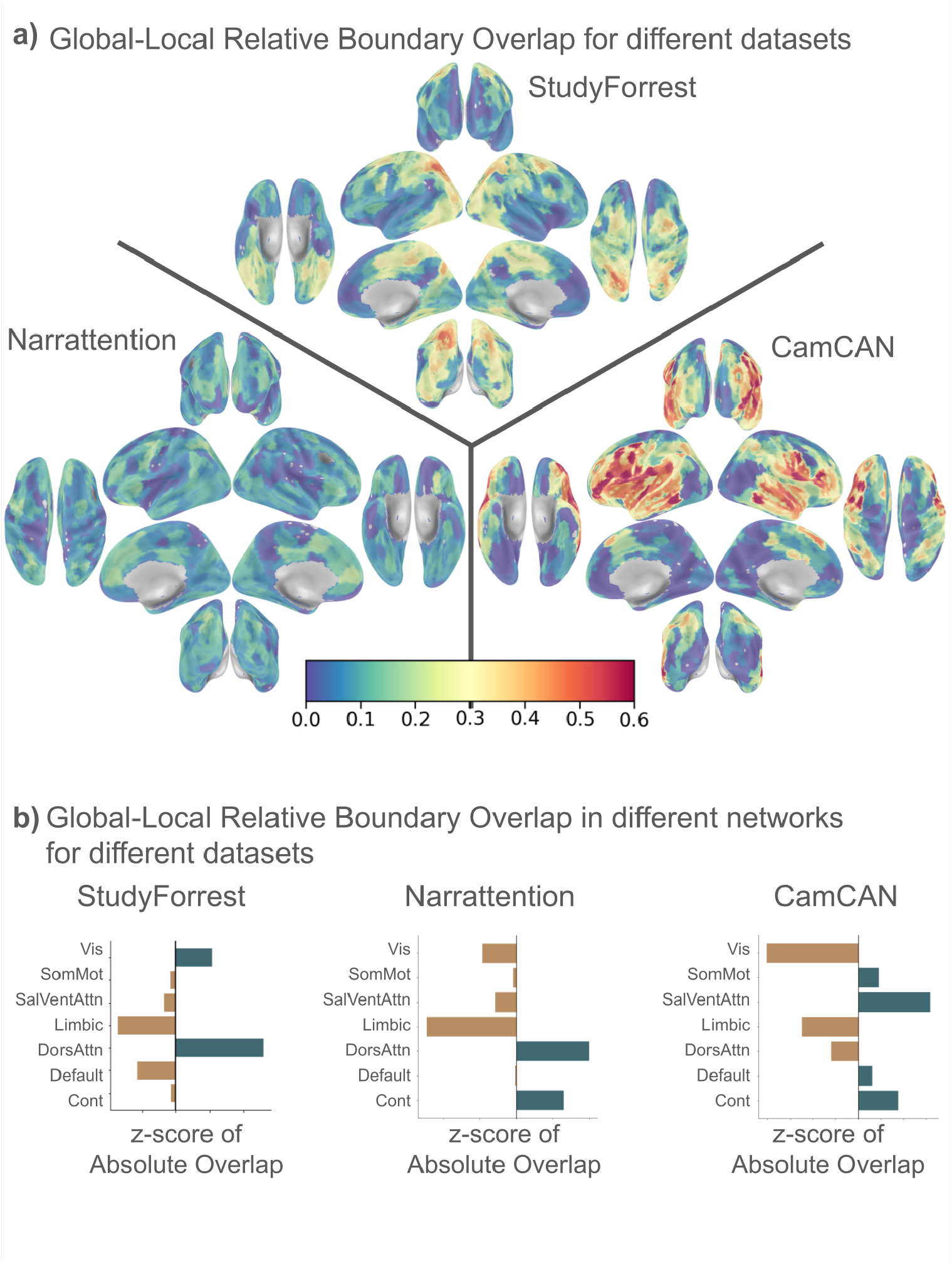
Global-Local state boundary overlaps (replication of 2) for all datasets. (a) Cortical maps showing the alignment between local ROI state transitions and the global brain state transitions. (b) Bar plots showing how the global-local alignment is distributed across functional networks.

## Appendix 6

### Appx.6: QPPs and Anti-correlated templates

We have translated the core QPP detection algorithm of the QPPLab software published by ***Xu et al. (2025***) from Matlab to Python. The QPP algorithm begins with an initial template pattern (which is chosen as a random location in data in the size of the window) and iteratively refines it through a convergence loop that continues until successive iterations achieve a correlation exceeding 0.9999. In each iteration, the function performs a sliding-window convolution between the current QPP template and the input data to generate a similarity time series, which is then z-score normalized and analysed using peak detection to identify timepoints where the data most closely matches the template. These peak locations are used to extract corresponding data segments, which are averaged to create an updated QPP template for the next iteration. This iterative refinement process allows the algorithm to bootstrap from an initial guess and converge on the most representative quasi-periodic spatiotemporal pattern present in the data, returning both the final QPP template and a normalized similarity time series indicating the temporal occurrences and strengths of the pattern throughout the dataset. Here the window size we use is 22 seconds. Figure1 shows the comparison between our template patterns and the QPPs for both whole-brain global data, and within-ROI local data. Overall, we see that both methods extract equivalent neural patterns. The correlation between the QPP pattern and our template pattern is close to r=1 at parts of the cycle and close to r=-1 at other parts of the cycle. QPPs have a fixed window size where both ends of our anti-correlated template pattern is expressed. The QPPs are defined on the whole brain scale, but you can see that they also are applicable at the local scale; and are still equivalent to one cycle of the anti-correlated template we define.

**Appendix 6—figure 1.**
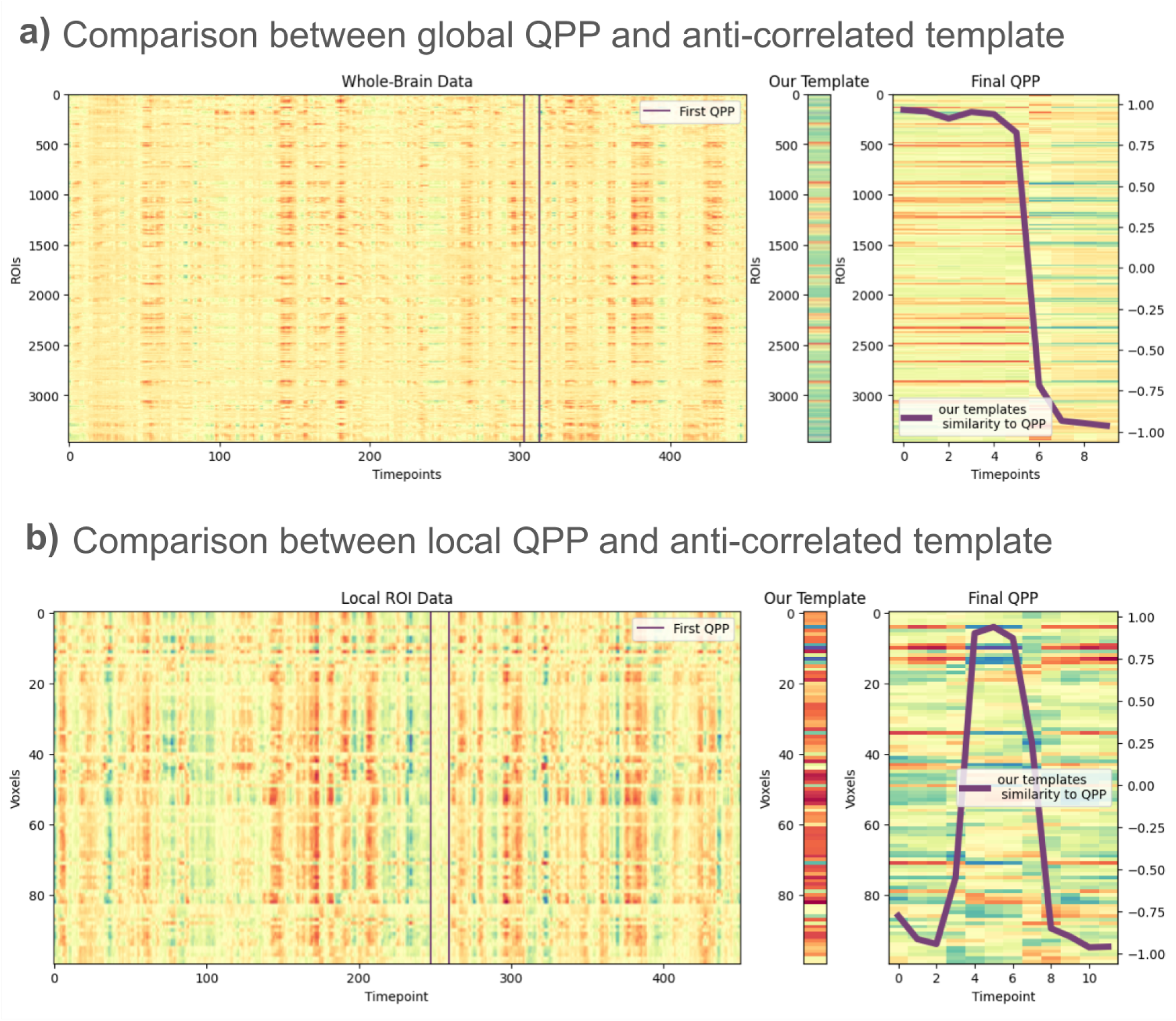
Comparing QPPs and anti-correlated templates. The anti-correlated patterns we uncover are almost identical to the QPPs defined by ***Xu et al. (2025***). QPPs constitute one cycle of the anti-correlated template we uncover (see both a and b, right). a) Left: ROI by time whole-brain activity. Middle: The anti-correlated template pattern. Right: The QPP, with the similarity of each timepoint of the QPP to our anti-correlated template. Similarity is quantified as Pearson r. b) Left: voxel by time local ROI activity. Middle: The anti-correlated template pattern. Right: The QPP, with the similarity of each timepoint of the QPP to our anti-correlated template. Similarity is quantified as Pearson r.

## Notes

https://github.com/drgzkr/Multi-Scale_Anti-Correlated

### Competing Interest Statement

The authors have declared no competing interest.

